# The price of a bit: energetic costs and the evolution of cellular signaling

**DOI:** 10.1101/2020.10.06.327700

**Authors:** Teng-Long Wang, Benjamin Kuznets-Speck, Joseph Broderick, Michael Hinczewski

## Abstract

Recent experiments have uncovered a fundamental information scale for cellular signaling networks: the correlation between input and output concentrations of molecules in a signaling pathway corresponds to at most 1-3 bits of mutual information. Our understanding of the physical constraints and evolutionary pressures that determine this scale remains incomplete. By focusing on a basic element of signaling pathways, the kinase-phosphatase enzymatic push-pull loop, we highlight the pivotal role played by energy resources available for signaling and their expenditure: the chemical potential energy of ATP hydrolysis, and the rate of ATP consumption. Scanning a broad range of reaction parameters based on enzymatic databases, we find that ATP chemical potentials in modern organisms are just above the threshold necessary to achieve empirical mutual information values. We also derive an analytical relation for the minimum ATP consumption required to maintain a certain signal fidelity across a range of input frequencies, where we quantify fidelity either through instantaneous or time-delayed mutual information. Attempting to increase signal fidelity beyond a few bits lowers the bandwidth, the maximum characteristic signal frequency that the network can handle at a given energy cost. The observed information scale thus represents a balancing act between fidelity and the ability to process fast-changing environmental signals. Our analytical relation defines a performance limit for kinase-phosphatase networks, and we find evidence that a component of the yeast osmotic shock pathway may be close to the optimality line. By quantifying the evolutionary pressures that operate on these networks, we argue that this is not a coincidence: natural selection on energy expenditures is capable of pushing signaling systems toward optimality, particularly in unicellular organisms. Our theoretical framework is directly verifiable using existing experimental techniques, and predicts that more examples of such optimality should exist in nature.

## I. INTRODUCTION

Survival for living cells depends in part on accurate and responsive signaling: the ability to collect enough information about the micro-environment to make decisions in response to external stimuli such as nutrients, hormones, and toxic agents [1]. This capacity to react to extracellular cues developed early in evolutionary history, and is now seen at all levels of biological organization, from chemotaxis in unicellular organisms [2–4] to the pathways that regulate cell differentiation and disease in multicellular life [5–8]. Despite the resulting diversity of biochemical networks that implement this signaling, information theory provides a powerful universal framework to quantify the amount of information transferred through a network, allowing comparisons between different systems [9].

Over the last decade a remarkable experimental consensus has emerged from such comparisons: studies of both prokaryotic and eukaryotic signaling pathways have found they can transmit at most *∼* 1 to 3 bits of information [10–17]. These values refer to mutual information (MI) between pathway input (concentrations of a molecule representing the signal) and output (concentrations of a downstream molecule produced by the network, sampled either at a single or multiple time points). MI is a measure of signal fidelity, representing the degree of correlation between input and output. Experiments have typically focused on a closely related quantity known as the channel capacity [18, 19]: the maximum MI achievable among all input distributions.

The consistently small channel capacities observed in cellular signaling pathways seem to indicate that cells operate with a fairly coarse representation of their surroundings: *n* bits of MI corresponds to being able to reliably distinguish between 2^*n*^ levels of the input, so a 1 bit pathway can only discriminate between “high” versus “low” signal concentrations. Though 1 bit is typical for MI measured at single time points, one can achieve higher MIs by focusing on output responses collected over several time points [14, 15], or by designing the experiment to isolate single-cell responses (as opposed to estimating MI from the responses of a population of cells) [17]. But these enhancements, which can push values to the 2-3 bit range, do not change the fundamental order of magnitude of the MI.

The central question we explore in this work is to what extent this fundamental information scale is shaped by the energy requirements of the underlying biochemical signaling networks. In order to transmit information, these networks necessarily need to operate out of equilibrium, fueled by processes like ATP hydrolysis that consume energetic resources. Recent research highlights these costs as an essential factor in understanding constraints on signaling [2–4, 20–23], often focusing on the ATP hydrolysis chemical potential difference 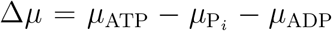 between the reactant (ATP) and products (ADP and inorganic phosphate, P_*i*_), quantifying the free energy available to drive the system per ATP. Crossing a certain minimum threshold of Δ*μ* is a prerequisite for a variety of signaling functions: accurate read-out of ligand-bound receptors [2, 3, 23], maintaining the phase coherence of oscillations in circadian clocks [20], or preserving the integrity of methylation-based “memory” to facilitate adaptation in chemotaxis [4]. This threshold is typically a few times larger (i.e. by a factor of *∼* 3 − 4 [2, 23]) than the energy scale of thermal fluctuations, *k*_*B*_*T*, where *k*_*B*_ is the Boltzmann constant and *T* the temperature. And indeed cells across the various domains of life maintain a sufficiently high Δ*μ ≈* 21 − 29 *k*_*B*_*T* [24] to enable such functions. The large value and remarkably narrow range of Δ*μ* observed in modern organisms opens up additional questions. The metabolic cycles that sustain Δ*μ*, constantly replenishing ATP as it is hydrolyzed, must almost necessarily have been far more inefficient and wasteful in the earliest stages of evolutionary history [25]. To what degree could organisms operating with smaller Δ*μ* still process information about their environment? What kinds of evolutionary pressures might have driven Δ*μ* to its modern range? And if the costs of individual signaling systems are non-trivial [2, 4], could natural selection have driven these networks toward optimized, energy-efficient solutions?

To investigate these issues, we focus on one of the canonical signaling circuits in biology, the kinase-phosphatase “push-pull loop”, which often forms a basic unit of more complicated signaling cascades [26–29]. An active kinase enzyme instigates the “push”, chemically modifying a substrate protein via phosphorylation (consuming ATP in the process), while a phosphatase enzyme provides the “pull”, dephosphorylating the modified substrate, reverting it to its original state. We derive the relationships between three facets of the system: i) the MI between the input (active kinase) and output (phosphorylated substrate) molecular populations; ii) the timescales over which the input signal varies; and iii) the energy requirements, expressed in terms of Δ*μ* and the rate of ATP consumption. Exploring the entire spectrum of kinase/phosphatase enzymatic parameters from bioinformatic databases, we find that physiological Δ*μ* values are just large enough to enable a MI of 1-2 bits for the widest possible parameter range. However to achieve this MI for signals that vary rapidly in time becomes more challenging, requiring both precise fine-tuning of parameters and a certain minimum rate of ATP consumption. In fact, taking advantage of results from optimal noise filter theory [30, 31], we derive a remarkably simple analytical relationship that describes the tradeoffs between minimum ATP rate, the MI, and the maximum characteristic signal frequency (the so-called bandwidth) which the push-pull network can handle. Verified via extensive numerical simulations across the whole gamut of enzymatic parameters, this relation is a novel theoretical prediction that can be directly tested in future experiments. We also demonstrate that analogous constraints hold even in the general case of time-delayed MI, where we look at correlation between the output and input signals separated by a time offset. The analytical relation rationalizes the observed range of MI by showing that values much higher than 1-2 bits would require sacrificing the ability to process fast-changing signals. Finally we explore the question of whether there exist evolutionary pressures that would push such a system to be energy efficient, optimizing the ATP consumption for a given target MI and bandwidth. Using a recently developed formalism relating metabolic costs to the strength of natural selection [32, 33], we show that these pressures can indeed be significant, particularly for single-celled organisms. We highlight a kinase-phosphatase loop in the yeast Hog1 signaling pathway as a system that may have been optimized by such pressures.

## II. THEORY

### A. Modeling an enzymatic push-pull loop

This push-pull network consists of two opposing reactions: a kinase enzyme instigates the “push”, chemically modifying a substrate protein via phosphorylation, while a phosphatase enzyme provides the “pull”, dephosphorylating the modified substrate, reverting it to its original state [26–29]. Since a single kinase can catalyze the phosphorylation of many substrate proteins, this loop can effectively act like an amplifier [28], translating a weaker signal (a small cellular population of an active kinase) into a stronger one (a large population of a phosphorylated substrate). Often the substrate itself is a kinase that can exist in catalyti-cally inactive and active states, with activation triggered by phosphorylation. In this case one can have multi-tiered signaling cascades enhancing the amplification (as shown schematically in Fig. 1A) with the active substrate produced by one loop serving as the kinase for a downstream loop [34]. More complex signaling networks are also possible, with multiple cascades connected by crosstalk through shared components [35], feedback from downstream to upstream populations [34], or activation requiring multisite phosphorylation [36]. However, the starting point for understanding any of these more complex signaling topologies is the behavior of a single loop, with a substrate activated / deactivated through a single phosphorylation site.

**FIG. 1.**
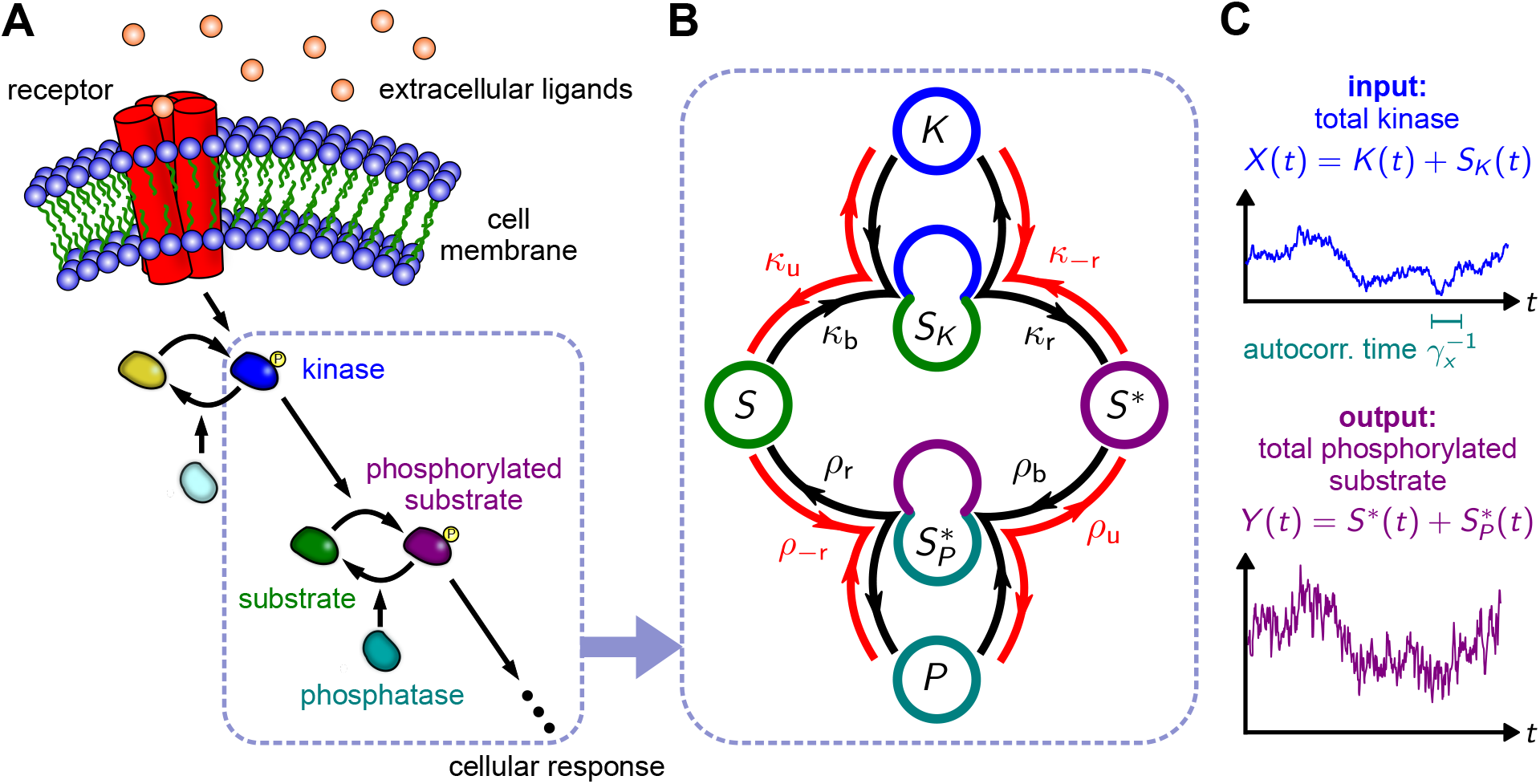
**(A)** A schematic signaling pathway involving cascades of kinase phosphorylation, initiated by a receptor embedded in the cell membrane that responds to extracellular ligands. The system we focus on will be one stage of the pathway, a kinase-phosphatase push-pull loop, highlighted in the dashed box. **(B)** The molecular species and reaction parameters of the push-pull loop. The kinase (*K*) binds to the substrate (*S*), forming the complex (*SK*) that catalyzes the production of phosphorylated substrate (*S\**). Phosphatase (*P*) binds to *S\**, forming a complex 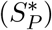 that catalyzes the dephosporylation of the substrate. Forward reaction / binding rates are labeled in black, while reverse reaction / unbinding rates are in red. **(C)** The loop serves to transduce an input signal, defined as the total population of kinase (bound or unbound), *X*(*t*) = *K*(*t*) + *SK*(*t*), into an output, defined as the total population of phosphorylated substrate, 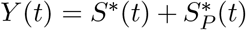. The input signal has a characteristic autocorrelation time 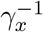.

The reaction scheme of a single push-pull loop is shown in Fig. 1B. Binding of free kinase (population *K*(*t*) at time *t*) to substrate (population *S*(*t*)) occurs with rate constant *κ*_*b*_, forming a kinase-substrate complex (population *S*_*K*_(*t*)). Phosphorylation of the substrate and its subsequent release constitutes the catalytic step, with rate *κ*_*r*_, yielding free phosphorylated substrates (population *S*^*\**^(*t*)). A phosphatase can subsequently bind, with rate *ρ*_*b*_, forming a phosphatase-substrate complex (population 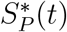), and catalyzing the dephosphorylation / release of the substrate with rate *ρ*_*r*_. These reactions also can occur in reverse: kinase-substrate unbinding (rate *κ*_*u*_), reverse kinase catalysis (rate *κ*_−*r*_), phosphatase-substrate unbinding (rate *ρ*_*u*_) and reverse phosphatase catalysis (rate *ρ*_−*r*_). Under physiological conditions some of these reverse rates may be negligible compared to their forward counterparts, but accounting for them is crucial to enforce thermodynamic consistency. In fact the product of the ratios of the reverse rates relative to the forward ones must satisfy a key thermodynamic relation arising from the principle of local detailed balance (closely related to the Haldane relation for enzymes) [37, 38],

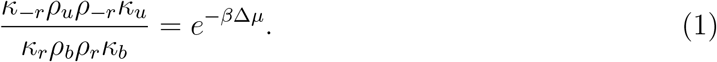

This relation is derived in the Supplementary Information (SI), and reflects the fact that for every complete traversal of the loop along the forward direction (clockwise along the black arrows in Fig. 1B) a single ATP molecule is removed from the environment, hydrolyzed, and the products ADP and inorganic phosphate P_i_ released back into the surroundings. Δ*μ* depends on the concentrations [ATP], [ADP], and [P_i_] through 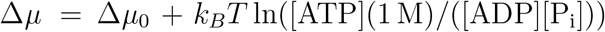, where 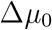 is the standard free energy of ATP hydrolysis (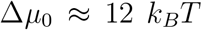 at room temperature [24]). Living systems expend energetic resources to maintain an imbalance of [ATP] relative to [ADP] and [P_i_], making Δ*μ* in physiological conditions larger than 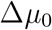. Despite the wide variety of metabolic pathways used to achieve this, measured Δ*μ* values in organisms from *E. coli* to humans lie within a relatively narrow range, Δ*μ ≈* 21 − 29 *k*_*B*_*T* [24]. This means reverse rates are sufficiently slow that the numerator in Eq. (1) is 9-12 orders of magnitude smaller than the denominator. One of the questions we tackle below is the significance of this disparity for transmitting information through the loop.

To quantitatively measure this information transfer, it is useful to explicitly describe the network behavior in terms of transducing an input signal into an amplified output, with degradation of the signal due to the stochastic nature of the reactions that mediate this process. We take the time-dependent input *X*(*t*) = *K*(*t*) + *S*_*K*_(*t*) to be the population of active kinases (both free and substrate-bound), and the corresponding output signal 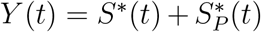 as the population of phosphorylated substrates (free and phosphatase-bound). For any specific system, the input kinases would be activated through a particular upstream signaling network. Here, however, we are interested in a more general problem: what is the effectiveness of this loop in processing a variety of possible input signals, spanning different amplitudes and timescales. The simplest mechanism that allows us to tune the dynamical characteristics of the input is to imagine the kinases activated at a constant rate *F* and deactivated at a constant rate *γ*_*K*_. We focus on the long-time limit where a stationary state has been achieved, and so *F* allows us to regulate the amplitude of the input signal while *γ*_*K*_ controls the autocorrelation time of the input fluctuations. While the analysis below could be done for other, system-specific models of the input, our choice allows us to explore a broad range of possible inputs to establish general bounds on information processing through the loop. With this input model, the reaction network model is fully specified. For a given set of parameters (drawn from distributions based on kinase/phosphatase biochemical information collected in enzymatic databases, as described below) we can derive analytical results for dynamical quantities using the linearized chemical Langevin approximation [39]. As shown in the SI, this provides excellent agreement with the exact kinetic Monte Carlo [40] simulation results in the parameter ranges of interest.

In focusing on how *X*(*t*) is transduced to *Y* (*t*), we frame our analysis in terms of three properties of the system. The first is the autocorrelation time of the input, 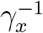, defined through 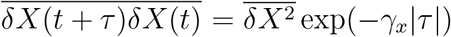, where the bar denotes an average over an ensemble of trajectories in the stationary state and 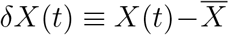. Note that instantaneous averages like 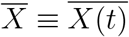 and 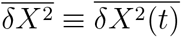 are independent of *t* in the stationary state. 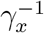is the characteristic timescale of the input fluctuations, and we will denote its inverse, *γ*_*x*_, as the effective “frequency” of the input. The second property is related to the mean rate at which phosphorylated substrates are produced through the catalytic reaction step, 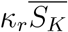, relative to the mean total number of activated kinases 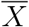. We define the gain parameter 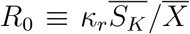 as a measure of the production of output for a given input level. Both *γ*_*x*_ and *R*_0_ can be expressed, to a good approximation, in terms of the reaction rates as follows (see SI for derivation):

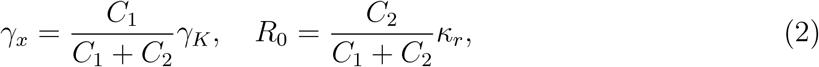

where 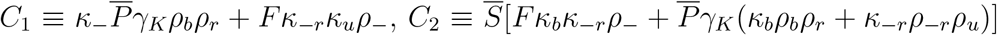. Here *κ*_−_ ≡ *κ*_*u*_ + *κ*_*r*_, *ρ*_−_ ≡*ρ*_*u*_ + *ρ*_*r*_. Note the dependence on mean unmodified substrate 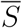 and free phosphatase 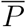 populations: these two numbers are free parameters that (along with the reaction rates) determine the network dynamics.

The final property of interest is the MI between input and output. Unless otherwise noted, we focus on the instantaneous stationary MI between *X*(*t*) and *Y* (*t*) at the same time point *t*, which we denote as ℐ. However in Sec. III C we also consider the MI between *X*(*t*) and *Y* (*t*+*α*), which we will denote with a subscript, ℐ _*α*_, where *α* is the time offset. This allows us to investigate whether including a time delay *α >* 0 (compensating for the finite propagation time of the signal) substantially changes the observed input-output correlation. For the instantaneous (*α* = 0) case, the MI is defined in terms of the joint probability *P* (*X, Y*) of observing input value *X* and output value *Y* at the same moment of time, and the corresponding marginal probabilities *P* (*X*) and *P* (*Y*),

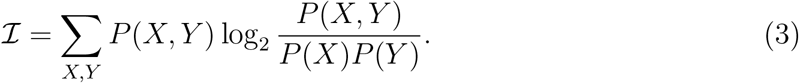

The value of *ℐ* is non-negative in all cases, and is measured in bits, with larger values translating to a greater degree of correlation between input and output. For our parameter ranges, *P* (*X, Y*) can be approximated as a bivariate Gaussian, and so we use an expression for *ℐ* valid in this limit that is more convenient to evaluate [19]:

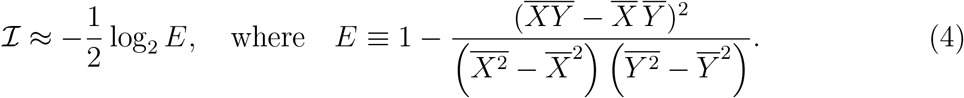

Here *E* = 1 − *ρ*^2^, where *ρ* is the Pearson correlation coefficient, and hence lies in the range 0 ≤ *E* ≤ 1. For *E* = 0 (or equivalently *ℐ* = *∞*) we have perfect correlation between the input and output signal, while *E* = 1 (*ℐ* = 0) corresponds to an output that is completely independent of the input. For the *α* ≠ 0 case, the expressions for *ℐ*_*α*_ would be analogous, except we use *P* _*α*_ (*X, Y*), the joint probability of observing *X* at time *t* and *Y* at time *t* + *α*. All averages, for example the cross-correlation 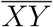, are then calculated with respect to *P* _*α*_ (*X,Y*).

### B. Determining the enzymatic parameter range

Once the input signal is specified through *F* and *γ*_*K*_, there are ten parameters related to the kinase, phosphatase, and substrate that determine the observables of interest *γ*_*x*_, *R*_0_, and *I* discussed above. These parameters are: 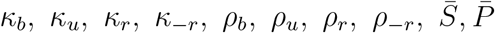. We know from surveys of enzymatic parameters that each of these quantities can span several orders of magnitude among different systems, often with an approximately log-normal distribution [41, 42]. To understand the performance limits of enzymatic loops in general, it makes sense to explore the entire range of biologically realistic parameters, rather than focus on a single choice of parameters. Existing online databases are excellent resources for this purpose, and Fig. 2 shows the resulting histograms of kinase / phosphatase parameters (full extraction details are available in the SI). For the substrate protein (which we take as a kinase) and the phosphatase, the concentrations [*S*] and [*P*] in Fig. 2A are derived from the PaxDb protein abundance database [43], using UnitProt gene ontology associations to identify kinases and phosphatases [44]. Enzymatic reaction parameters are available in the Sabio-RK database [45]. The reaction rates *κ*_*r*_ and *ρ*_*r*_ (Fig. 2D) are typically listed directly, but the others are most often in specific combinations: the Michaelis constants 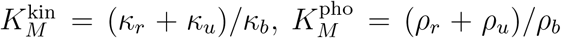 for kinase/phosphatase respectively (Fig. 2B) and the specificity ratios 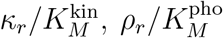 (Fig. 2C). For all of these parameters there is a paucity of data on phosphatases relative to kinases, but the phosphatase ranges seem to largely overlap with those of kinases. Thus for simplicity we take kinase and phosphatase parameters to have the same distributions (log-normal) and use a numerical fitting procedure to find an overall log-normal joint probability distribution for the eight underlying model parameters represented in the data: 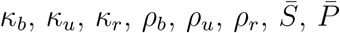 (see SI). Note that data in concentration units (like [*S*] and [*P*] in molars) is converted to mean abundance (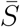 and 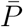) by assuming a volume of 30 fL (comparable to the cytoplasmic volume of yeast [24, 46]). This procedure is designed so that the resulting joint distribution yields marginal probability densities (solid curves in Fig. 2) that exhibit good agreement with the histogram data for any of the measured parameter combinations. Despite this agreement, we note that the joint distribution likely spans a portion of the parameter space larger than the true distribution of biological values: this is because it cannot fully capture correlations between different parameters. (Such correlations are difficult to reconstruct since many database entries are incomplete, containing some but not all of the enzymatic parameters.) For our purposes, having a distribution that effectively acts like a superset of the biological distribution is fine: whatever performance bounds we infer from the whole distribution will then also apply to the subset of the distribution that corresponds to current real-world systems. Moreover this also allows us to explore a larger enzymatic design space, which may have been accessible at earlier points in evolutionary history.

**FIG. 2.**
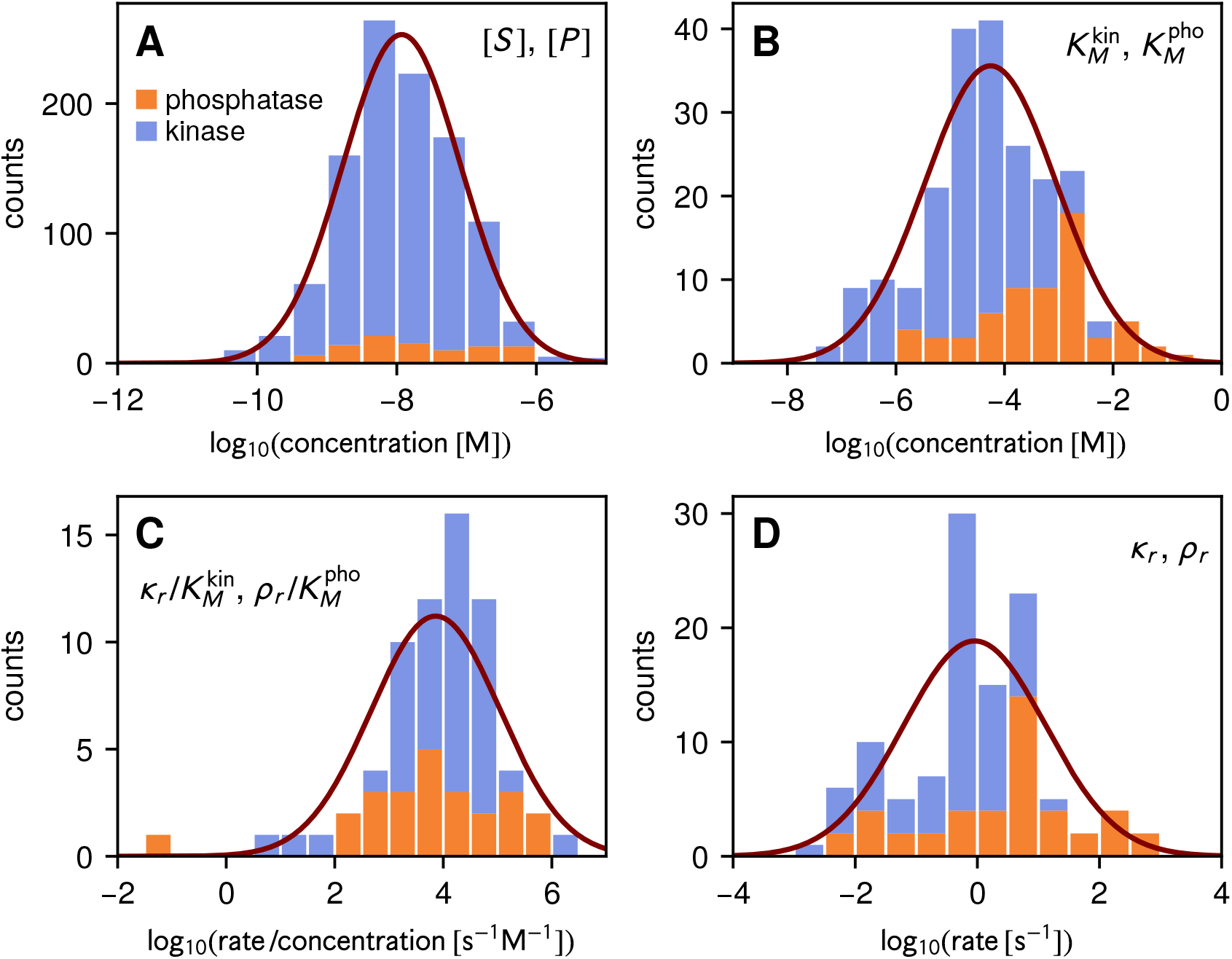
Enzymatic parameter ranges for kinases/phosphatases based on the PaxDb [43] and Sabio-RK [45] databases. Because of the relative lack of phosphatase data (orange histograms) relative to kinases (blue histograms), we fit an overall log-normal joint probability to the total data set including both kinases and phosphatases. The marginal distributions from that global fit are plotted as purple curves. The parameters are as follows: **(A)** kinase substrate [*S*] and phosphatase [*P*] concentrations; **(B)** kinase/phosphatase Michaelis constants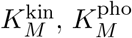; **(C)** the corresponding specificity ratios 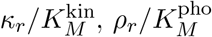; **(D)** kinase/phosphatase catalytic rates *?*_*r*_ and *?*_*r*_.

Two of the model parameters are still unaccounted for: the reverse reaction rates *κ*_−*r*_ and *ρ*_−*r*_. Though usually small in magnitude and typically not measured in enzyme kinetic assays, we also know that they are crucially related to Δ*μ* through the local detailed balance relation of Eq. (1). Thus, as explained in the next section, these become important free parameters that we can vary to explore signaling efficiency and its dependence on Δ*μ*.

## III. RESULTS

### A. Minimum cost of transmitting information

Given the model described above, with a parameter set drawn at random from the empirical joint distribution, we can ask a basic first question: what is the minimum chemical potential difference Δ*μ* required to achieve a certain mutual information *ℐ*? The answer will depend on the nature of the input signal *X*(*t*), and thus we would like to test different effective input frequencies *γ*_*x*_. To do this we will fix the mean free kinase concentration at the level of a low amplitude input, [*K*] = 5 nM, and vary *γ*_*K*_, which varies *γ*_*x*_ according to Eq. (2) with 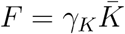 for fixed 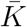. In the SI we also show the same analysis for [*K*] = 0.5 and 50 nM, with results qualitatively similar to those described below. After drawing enzyme parameters from the joint distribution and specifying *γ*_*x*_ at a given [*K*], the only two free parameters are the reverse reaction rates *κ*_−*r*_ and *ρ*_−*r*_.

Fig. 3A shows a contour diagram of *ℐ* as a function of *κ*_−*r*_ and *ρ*_−*r*_ for a sample enzyme parameter set and value of *γ*_*x*_. Superimposed are dotted lines of constant Δ*μ* from Eq. (1). If one were interested in achieving a particular *ℐ* value, for example *ℐ* = 1 bit, one can then numerically determine the *κ*_−*r*_ and *ρ*_−*r*_ point along the *ℐ* = 1 bit contour where Δ*μ* is smallest. For this specific enzyme parameter set and *γ*_*x*_, the value turns out to be Δ*μ* = 6.72 *k*_*B*_*T*, which would then be recorded as the minimum necessary Δ*μ* to achieve 1 bit of MI. Note that it is not guaranteed that a minimum Δ*μ* solution exists for every parameter set sampled from the joint distribution. If the *ℐ* contours plateau at a maximum less than 1 bit, no possible Δ*μ* will allow that particular system to achieve the desired MI target. We will return to this important point below.

**FIG. 3.**
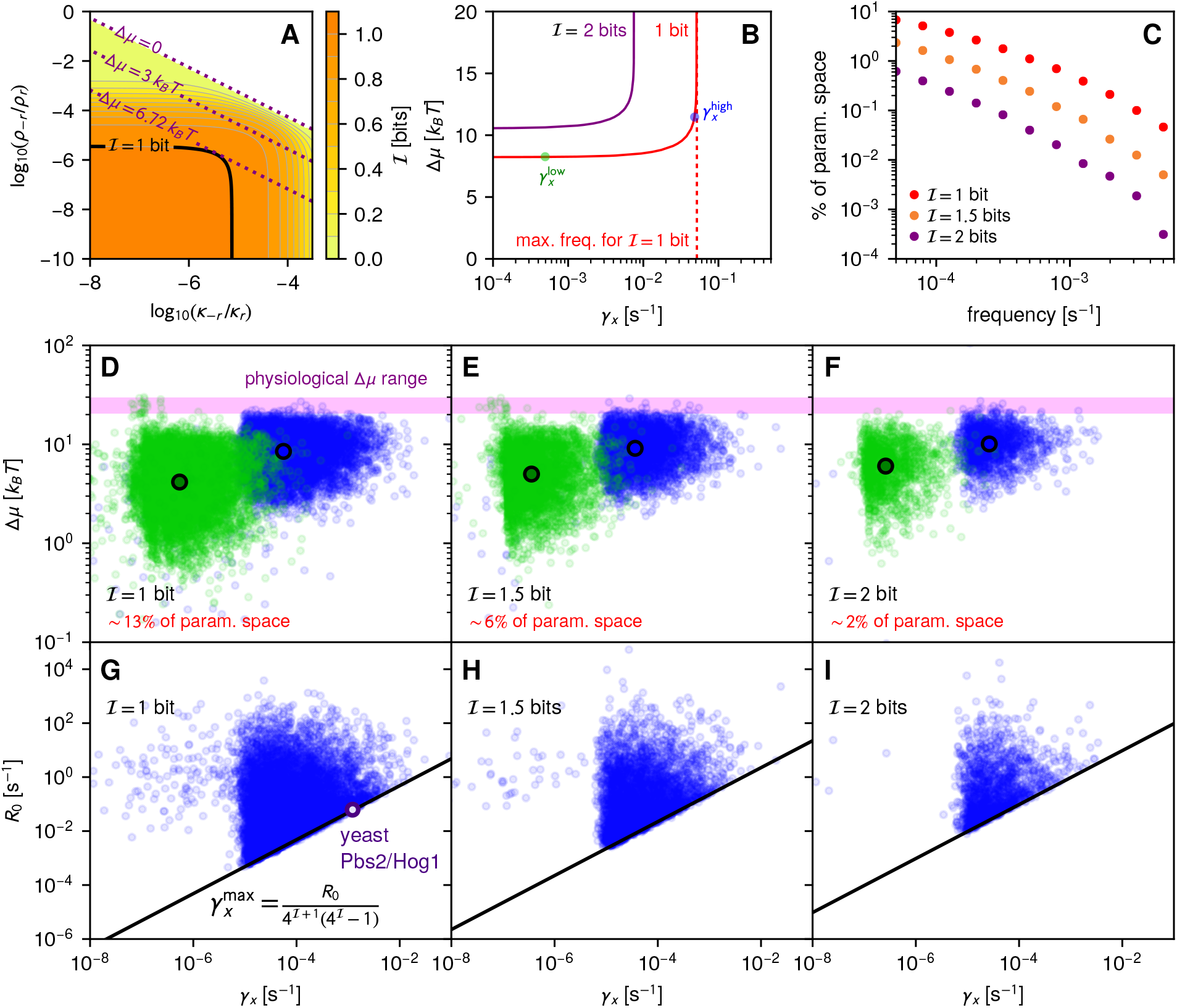
**(A)** A representative contour diagram of *ℐ* (solid curves) as a function of *κ*_−*r*_ and *ρ*_−*r*_ for a parameter set drawn randomly from the joint distribution. Dotted lines denote contours of constant Δ*μ*. In this case Δ*μ* = 6.72 *k*_*B*_*T* is the smallest value at which the system can achieve *ℐ* = 1 bit. **(B)** For a sample parameter set, the minimum Δ*μ* needed to achieve *ℐ* = 1, 2 bits as a function of input frequency *γ*_*x*_. For the 1 bit case, the dashed line represents 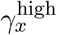, the maximum *γ*_*x*_ compatible with ℐ = 1 bit for this parameter set. As described in the text, we highlight two points along the curve: one at a frequency 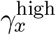 at roughly 95% of the bandwidth, and the other at frequency 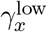 at roughly 1% of the bandwidth. The points will be plotted for a many random draws of the enzyme parameters from the joint distribution in the lower panels of the figue. **(C)** For each target value of ℐ= 1, 1.5, 2 bits, the percentage probability of randomly drawing a parameter set that has a 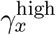 higher than a given frequency. **(D-F)** The distribution of 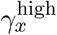 (blue) and 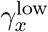 (green) for many random parameter draws, keeping only those that can achieve *ℐ* = 1 bit (D), 1.5 bits (E), or 2 bits (F). The probabilities of successfully drawing such a set are shown in red in each panel. The blue and green circles denote the median of each distribution respectively. **(G-I)** The same 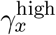 distributions as in panels (D-F), except plotted in terms of gain *R*_0_ on the vertical axis. The solid line is the analytical maximum bandwidth bound 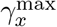 of Eq. (5). The purple circle in panel G shows the estimated result for the near-optimal yeast Pbs2/Hog1 system.

If one keeps the enzyme parameters (other than *κ*_−*r*_ and *ρ*_−*r*_) fixed, and just varies *γ*_*x*_, an interesting trend appears in the minimum Δ*μ* results. Fig. 3B shows two examples of minimum Δ*μ* curves, for target *ℐ* values of 1 and 2 bits respectively. For a given *ℐ* target, the minimum Δ*μ* is nearly constant at low input frequencies, but then increases rapidly and diverges at a maximum frequency which we will dub the “bandwidth” of the system. This intuitively makes sense: the higher the input frequency, the more rapid the catalytic reaction rates needed to accurately transmit the signal through the system, increasing the required Δ*μ* threshold. However, there is an inherent limit, given finite enzyme catalysis rates. Above the bandwidth, whose value depends on the enzyme parameters, the system can no longer achieve the target *ℐ*. The higher the informational burden (i.e. increasing the target *ℐ* from 1 to 2 bits) the lower the bandwidth: if one desires higher fidelity transmission, the range of transmissible signal frequencies will suffer.

To make more sense of these results, it is useful to look at a broad sample of enzyme parameters rather than a single set. To visualize global behaviors, we will calculate two numerical results for each set drawn from our joint distribution. The procedure is as follows: i) Sample an enzyme parameter set from the distribution; ii) Determine if it can achieve our target *ℐ* for any input frequency; iii) If the answer is yes, vary *γ*_*K*_ until one finds the maximum possible value 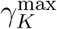 where one can still achieve the *ℐ* target. iv) Calculate the minimum Δ*μ* for an input signal very near the bandwidth frequency, where 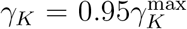. We will call this result Δ*μ*^high^. The corresponding input frequency is 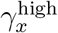. v) Analogously, calculate the minimum Δ*μ* for an input signal with a frequency much lower than the bandwidth, where 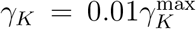. This set of results we denote as Δ*μ*^low^ and 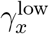. Fig. 3B shows the two points 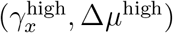 and 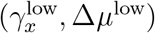) as blue and green dots respectively for that particular parameter set at *ℐ* = 1 bit. These two points encapsulate several key features of the minimum Δ*μ* versus *γ*_*x*_ curve: Δ*μ*^low^ roughly corresponds to an “entry level” price, the minimum ATP hydrolysis chemical potential necessary to transmit the signal at any frequency, while the difference Δ*μ*^high^ − Δ*μ*^low^ is the premium one has to pay to transmit signals near the highest possible frequencies. The value 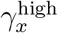 approximately corresponds to the bandwidth.

If one were to make numerous draws from the parameter distribution, and plot 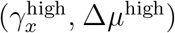 and 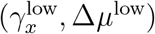) for each draw, one would get a cloud of blue and green dots. These are shown in Fig. 3D-F for target *ℐ* of 1, 1.5, and 2 bits respectively. As mentioned above, not every draw will lead to a parameter set that can achieve the target, and the plots are labeled by the fraction of draws that are capable of reaching that particular value of *ℐ*. That fraction decreases with *ℐ*, from 13% for *ℐ* = 1 bit down to only 2% for *ℐ* = 2 bits. As *ℐ* increases not only does it become progressively more difficult to find enzymatic parameters compatible with higher fidelity, but the accessible frequency range becomes more restricted. Fig. 3C shows the percentage of the parameter space that can achieve bandwidths higher than a given frequency for different *ℐ*. For example let us consider the frequency 1.22 *×* 10^−3^ s^−1^, which is the *ℐ* = 1 bit bandwidth for the yeast Pbs2/Hog1 system described in detail in Sec. IIIC. (This system is part of the Hog1 osmotic stress response pathway, whose overall bandwidth has been experimentally estimated to be of a similar scale [47]). From Fig. 3C it is evident that only about 0.41% of the draws from the parameter distribution have 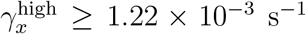 for a target *ℐ* = 1 bit. If one were to attempt to transmit signals at such high frequencies for *ℐ* = 2 bits, the fraction of compatible parameter space shrinks to a miniscule 9 *×* 10^−3^%. This reflects the exquisite fine-tuning required to put together a set of enzymatic loops capable of responding to quick, life-or-death variations of the external environment on time scales of a couple of minutes. Going much beyond *ℐ* = 1 bit and maintaining fast response times for a single push-pull loop is extremely difficult, and hence it makes sense that biology settles for *ℐ* in the vicinity of 1 bit in many circumstances.

Going much below 1 bit poses another set of difficulties, since such systems would not even be able to reliably transmit the difference between high and low values of input signal. For signaling that can occur over longer timescales (hours instead of minutes) it becomes much easier to find compatible parameter sets, with the median of the distribution of 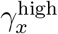 for *ℐ* = 1 bit around *∼* 6 *×* 10^−5^ s^−1^.

From the perspective of costs, the bulk of the distribution of entry level prices Δ*μ*^low^ for *ℐ* = 1 bit is ≳1 *k*_*B*_*T*. Any system much below this would be too close to equilibrium (reverse rates comparable to forward rates) for effective information transfer to occur. The median of the Δ*μ*^low^ distribution in Fig. 3C is 4 *k*_*B*_*T*, increasing to about 6 *k*_*B*_*T* for *ℐ* = 2 bits in Fig. 3E. These values are on the same scale as estimates of minimum Δ*μ ∼* 4 *k*_*B*_*T* ln 2 required for 99% accurate readout of a ligand-bound receptor via the activation of a downstream molecule, assuming an arbitrarily slow readout process [23]. In that system (as in ours), processing information at faster time scales requires large Δ*μ*. Indeed we find that the median values for Δ*μ*^high^ range between 8 − 10 *k*_*B*_*T* for *ℐ* = 1 − 2 bits. The minimum Δ*μ* near the bandwidth is typically shifted up by about 4 *k*_*B*_*T*, reflecting the premium necessary to transmit near the frequency limit. Paying this premium is worthwhile: frequencies 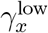 accessible at Δ*μ*^low^ prices are likely far too low to have biological relevance, with the distributions of 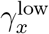 largely below 10^−5^ s^−1^. To get the ability to respond to signals at more biologically reasonable time scales thus means being capable of transmitting closer to the bandwidth, making Δ*μ*^high^ a more useful measure of minimum biological costs.

The Δ*μ*^high^ distributions show that it is possible to have signaling systems that transmit at least 1 bit of MI and operate at Δ*μ* lower than the current physiological range (Δ*μ ≈* 21−29 *k*_*B*_*T* [24], indicated in pink in Fig. 3D-F). This is true even for systems with the fastest responses (large 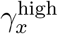 near the right edges of the distribution). This means that one can imagine enzymatic signaling systems in the earliest stages of evolutionary history that can reliably distinguish high and low inputs even before ATP metabolism (maintaining high ATP concentrations relative to ADP and P_*i*_) reached its modern levels of efficiency.

In fact a fascinating universal feature of the distributions is that the physiological Δ*μ* range lies just above the top edge of the distributions. Naively it would seem as if the physiological values are just high enough to allow these signaling loops to transmit *ℐ* = 1 *–* 2 bits across the broadest possible parameter subset. This gives evolution the largest possible space in which to tweak tradeoffs between fidelity and response times without running into chemical potential limitations. Of course Δ*μ* influences not just signaling networks but the entire range of cellular functions, so it is impossible to say with certainty what factors played the largest role in determining the values of Δ*μ* we see in present-day organisms. But at least from the perspective of signaling at the level of a push-pull loop, it is clear that Δ*μ ≈* 21 − 29 *k*_*B*_*T* is more than good enough for basic information transfer needs, and there would be no benefit in having a system with substantially higher Δ*μ*. To maintain Δ*μ* = 40 or 50 *k*_*B*_*T* for example, would require significant additional metabolic resources, with little payoff in terms of either *ℐ* or bandwidth.

### B. Analytical bound describes tradeoff between bandwidth and information

The results above already illustrated the tradeoff between bandwidth and MI, with parameter sets that achieve very large 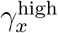 becoming progressively harder to find as the target *ℐ* increases. Can we understand this relationship in more detail? For this purpose we take advantage of optimal noise filter theories, originally developed in the context of signal processing [48–50], and in recent years applied to a variety of biological signaling networks [30, 31, 51–54]. The original motivation involved designing a filter for a signal corrupted by noise, such that the output matched the uncorrupted input signal as closely as possible. In the biological context, this same framework allows us to put bounds on the maximum MI achievable between input and output signals for given input and enzymatic parameters. As shown in the SI, our enzymatic push-pull loop can be approximately mapped onto an effective two-species input-output system, which is then amenable to analytical treatment using the Wiener-Kolmogorov optimal filter theory [30, 48–50].

The end result is a remarkably simple analytical relation between the maximum possible bandwidth 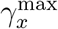 achievable given a target value of *ℐ*,

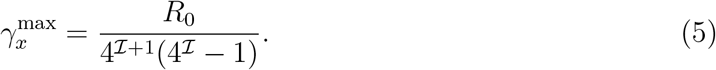

The only other enzymatic parameter that appears in the relation is the gain *R*_0_, a measure of output production relative to the input. Fig. 3G-I shows the same parameter set distribution as the 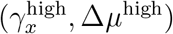 points in Fig. 3D-F, except replotted in terms of 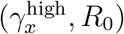, where *R*_0_ is the gain for each parameter set. The solid line is the bound of Eq. (5). Even though this bound is based on an approximation of the full enzymatic system, and hence is not guaranteed to be exact, it still provides an excellent cutoff for the distribution of 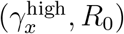 points. For systems at a certain *R*_0_, we see that as *ℐ* is increased and the denominator in Eq. (5) gets larger, the maximum bandwidth 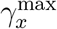 shifts to lower values. If we are interested in a fast response time, increasing *ℐ* systematically reduces the compatible parameter space, since we are forced to rely on cases with larger and larger *R*_0_. Thus Eq. (5) rationalizes the earlier observation of limited options for networks that can simultaneously respond to signals fluctuating on minute time scales and achieve *ℐ* significantly larger than 1 bit.

### C. Time-delayed mutual information

Up to now we have focused on the instantaneous MI *ℐ* between input and output, which is constrained not just by noise in signal transduction, but also by the fact that changes in the input take a finite time to propagate into the output. We can compensate for the finite propagation speed by using the time-delayed MI *ℐ*_*α*_, which describes the correlation between the input *X*(*t*) and the output *Y* (*t*+ *α*) at a later time [55, 56]. Indeed some of the gains in MI observed in experiments collecting outputs over multiple time points [14, 15] reflect the fact that later output time points can have higher correlations with the current input. Fig. 4A shows the typical behavior of *I*_*α*_ versus *α*, for a parameter set that achieves instantaneous MI *ℐ*_0_ = 1 bit at *α* = 0, with 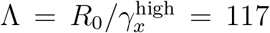. The MI reaches a maximum value 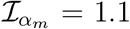 bits at time delay *α*_*m*_ *>* 0, which is usually a fraction of the characteristic time scale 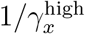. The results for the chemical Langevin approach (solid curve) are consistent with kinetic Monte Carlo simulations (circles). For completeness we also show the behavior of *ℐ*_*α*_ for *α <* 0, which describes the extent to which the output can anticipate future input signals. As expected, *ℐ*_*α*_ < *ℐ*_0_ for *α <* 0, and *ℐ*_*α*_ decreases as the time offset | *α* | becomes larger, reflecting the difficulty of forecasting the input as one looks further into the future.

The approximate two species Wiener-Kolmogorov optimal filter theory described in the previous section easily generalizes to incorporate time delay [54] (details in the SI). It yields an analytical bound *I* ^max^(Λ) on the MI regardless of time offset *α*:

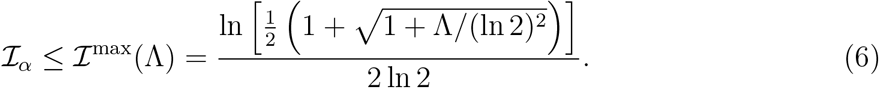

**FIG. 4.**
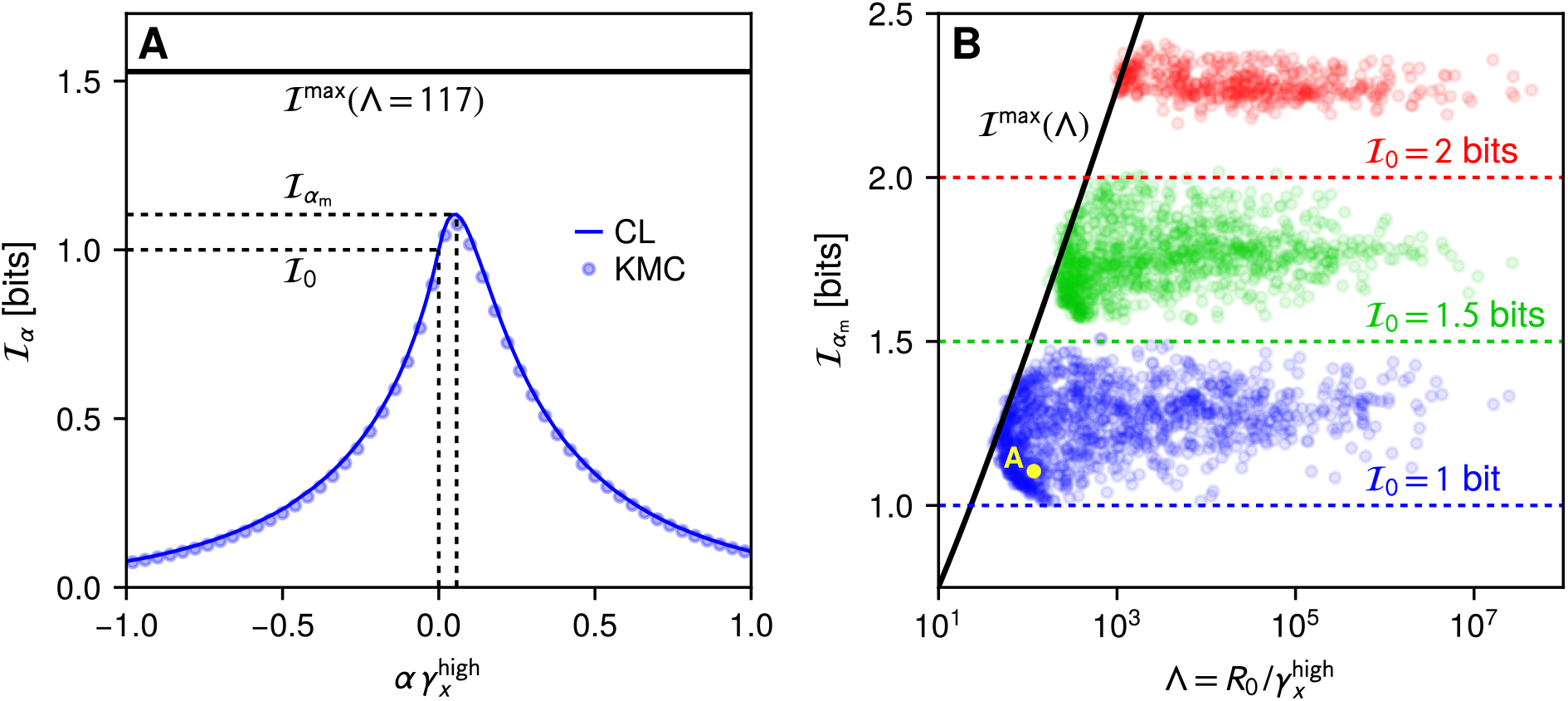
**(A)** *ℐ*_*α*_, the MI between input and output with a time offset *α*, for a sample parameter set where the system achieves instantaneous MI *ℐ*_0_ = 1 bit. The maximum 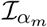 is achieved at a finite time delay *α*_*m*_ *>* 0. Results from the chemical Langevin approximation (CL) agree with kinetic Monte Carlo simulations (KMC). The analytical bound *ℐ*^max^(Λ) from Eq. (6) is shown as a horizontal line, with Λ = 117 for this parameter set. **(B)** The three sets of points correspond to the parameter sets distributions used in Fig. 3G-I, with *ℐ*_0_ = 1 bit (blue), 1.5 bits (green), and 2 bits (red). For each parameter set, the maximum time-delayed 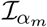 is plotted versus 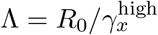. The bound *ℐ*^max^ (Λ), based on the approximate two species optimal filter theory, is shown as a black curve.

The value *I* ^max^(Λ) = 1.53 for Λ = 117 is shown as a horizontal line in Fig. 4A, so for this parameter set the actual maximum 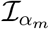is not near the bound. However there are systems that can achieve *I*^max^(Λ). Fig. 4B shows parameter sets from Fig. 3G (blue), 3H (green), and 3I (red), corresponding to systems with instantaneous MI values of *ℐ*_0_ = 1, 1.5, or 2 bits respectively. For each parameter set, the maximum time-delayed 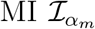 is plotted versus 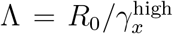. The bound *I*^max^(Λ) is plotted as a solid curve, and there is a subset of systems with performance near the bound. As we noted earlier, the two species optimal filter theory is an approximation for the full enzymatic system, and hence the bound does not have to be exact. But it still provides a close estimate of the MI limits for the full system, with the largest violations of the bound only *∼* 4%. The benefits of time delay are particularly evident for parameter sets with lower *ℐ*_0_: some cases with *ℐ*_0_ = 1 bit exhibit 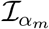approaching 1.5 bits.

### D. Optimality and the yeast Pbs2/Hog1 push-pull loop

Returning to the discussion of instantaneous MI, there is an alternative way of thinking about the *R*_0_ versus 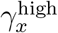 results in Fig. 3G-I. Imagine a system working at 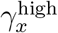 with a certain gain parameter *R*_0_ and achieving a target value *ℐ*. Comparing other parameter sets with the same bandwidth 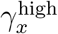 and target *ℐ* (taking a vertical slice of one of the panels in Fig. 3G-I), they will have a variety of different *R*_0_ values, but all of these will be bounded from below by the minimum value

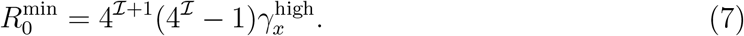

When 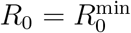, the system sits on the optimality line of Eq. (5), with 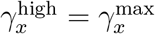.

The discrepancy between *R*_0_ and 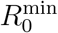 for a given system allows us to see how close the signaling behavior is to optimality. Let us take a concrete biological example: the Pbs2/Hog1 enzymatic push-pull loop from yeast, part of the Hog1 signaling pathway that allows the organism to respond to osmotic stress. As described in the SI, key parameters for this system can be estimated based on an earlier model [46] fit to microfluidic experimental data where yeast was exposed to periodic salt shocks [57]. The results for the bandwidth and gain for *ℐ* = 1 bit are: 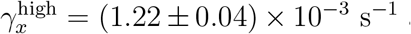 and *R*_0_ = 0.0621 *±* 0.0001 s^−1^, with the error bars reflecting uncertainties due to unknown parameters (where we used priors based on the log-normal distributions in Fig. 2.) The scale of the predicted bandwidth 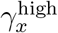 is consistent with microfluidic estimates. Ref. [47] found a steep dropoff in the mean amplitude of the Hog1 response to periodic step-like changes in external osmolyte concentrations when the frequencies of the changes increased from 10^−3^ s^−1^ to 10^−2^ s^−1^. At frequencies beyond the dropoff the Hog1 output can no longer reproduce the osmolyte input at high fidelity. Though the form of the input in this case is different than in our model, and the experiment probes the entire pathway rather than just the Pbs2/Hog1 component, the similarity in scales to our 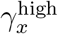 value suggests that the Pbs2/Hog1 system may play a major role in determining the bandwidth of the whole pathway (since the bandwidth of the whole is constrained by the bandwidths of the components).

Intriguingly, the estimated gain *R*_0_ is very close to the minimum possible value 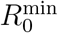 for signaling at the bandwidth 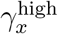 with *ℐ* = 1 bit, as seen in Fig. 3G. Using Eq. (7), we find 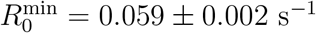. This naturally leads to the question: is the fact that this system lies so close to optimality a coincidence, or are there reasons why natural selection might favor minimizing *R*_0_ in this case? To answer this question, we first have to consider the relationship between gain and ATP consumption.

### E. Minimum ATP consumption to achieve a certain signaling fidelity and bandwidth

This bound on the gain parameter in Eq. (7) is directly related to the metabolic cost of signaling, since higher production of the output per given input level will generally require a higher rate of phosphorylation events. We can roughly quantify the average rate of phosphorylation: in the stationary state this is just the mean rate of the kinase-catalyzed reaction step, 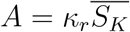. Assuming one ATP hydrolyzed per reaction, *A* is the mean rate at which ATP is consumed by the system, and is related to *R*_0_ through 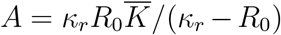, as shown in the SI. In the enzymatic parameter ranges we consider, *κ*_*r*_ is typically much larger than *R*_0_, so we can approximate this relation as 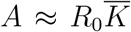. Using Eq. (7) we can then estimate the minimum possible ATP consumption rate given a target *ℐ* and bandwidth 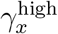:

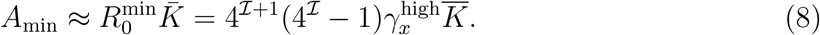

Fig. 5A shows the same parameter set values as the 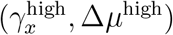 points in Fig. 3D for *ℐ* = 1 bit, except plotted in terms of 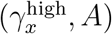. The *A* values are exact, but the approximate relation of Eq. (8) provides an excellent lower bound on the distribution. Qualitatively, the individual elements of Eq. (8) all make intuitive sense. An increase in any of the con-stituent factors (the mean free input kinase population 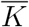, the target information *ℐ*, the bandwidth 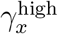) puts greater demands on the signaling system, requiring more catalytic activity and hence faster ATP consumption. Note that the above results are easily generalized if the reaction step consumes more than one ATP: for example the effective model for yeast Pbs2/Hog1 discussed above involves phosphorylation at two sites, which would lead to expressions for *A* and *A*_min_ getting a prefactor of two.

**FIG. 5.**
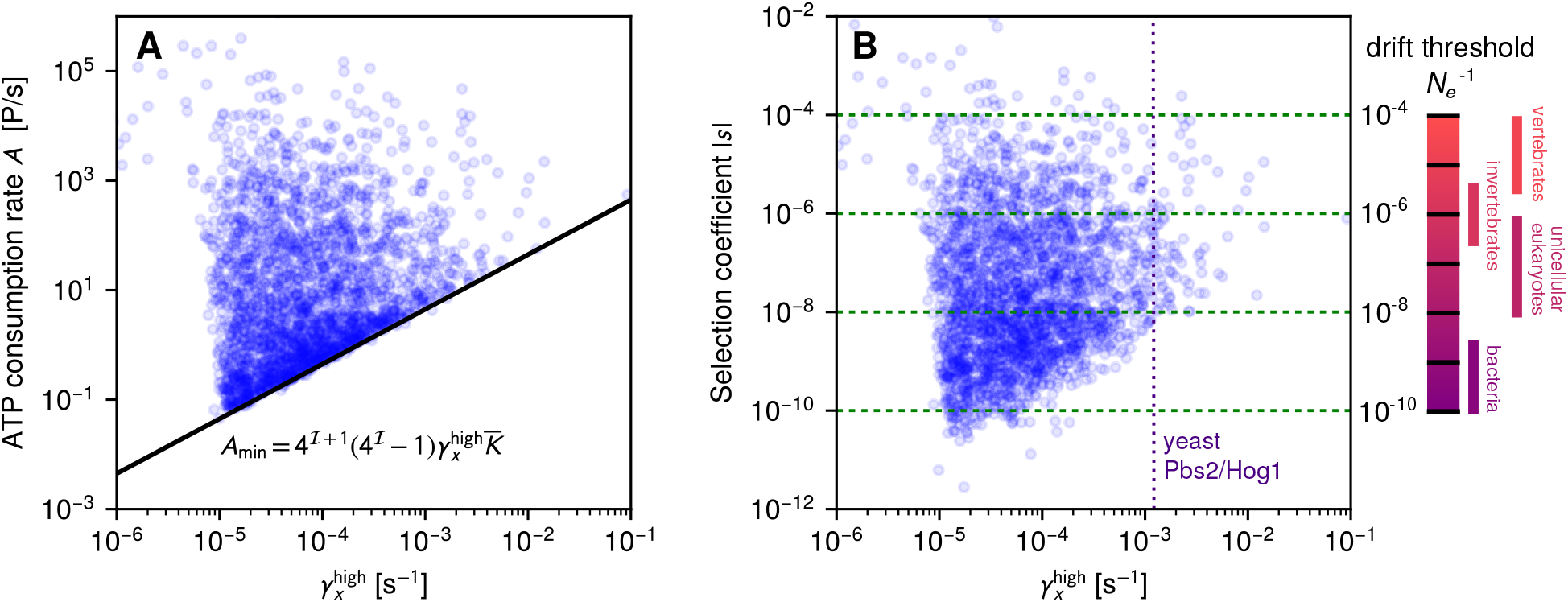
**(A)** The same 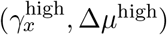 point distribution as in Fig. 3C for *ℐ* = 1 bit, except plotted in terms of ATP consumption rate *A* on the vertical axis. The solid line is the approximate lower bound *A*_min_ on ATP consumption given by Eq. (8). **(B)** This distribution replotted with selection coefficient |*s*| on the vertical axis. |*s*| quantifies the fitness cost associated with a system that achieves the target *ℐ* = 1 bit but is sub-optimal in ATP consumption, relative to an optimal variant where *A* = *A*_min_. The value of |*s*| becomes evolutionarily significant when it is higher than a “drift threshold” 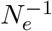, where *N*_*e*_ is the effective population of the organism (a measure of genetic diversity). The ranges of 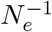 for different classes of organisms are shown on the right [32, 58]. The vertical dotted line corresponds to the estimated 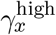 for the yeast Pbs2/Hog1 system.

### F. Evolutionary pressure on the metabolic costs of signaling

It is clear from Fig. 5A that for many parameter set choices the ATP consumption rate *A* is significantly larger than for a system near optimality (*A ≈ A*_min_) given the same *ℐ* and 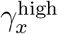. Let us consider a specific scenario where the bandwidth 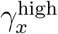 and the target *ℐ* are sufficient for the biological function of the signaling i.e. there are rapidly diminishing fitness returns in going to higher bandwidth and signal fidelity. In this scenario a system with *A > A*_min_ has no significant adaptive advantage over one with *A ≈ A*_min_, but instead incurs a fitness penalty because of the superfluous ATP consumption. Would there be evolutionary pressure on this sub-optimal system to move toward optimality?

The answer to this question has practical ramifications, because it will allow us to predict whether we should expect to see natural enzymatic push-pull loops cluster around the optimality line (as we saw in the yeast Pbs2-Hog1 example). The alternative, in the absence of strong evolutionary pressure to optimize, is a wider dispersion, more similar to Fig. 5A where the points are drawn at random from the enzymatic parameter distribution. Note that this is a question that is directly amenable to future kinetic experiments: for systems where we can fully characterize the enzymatic parameters of the push-pull loop (for both the kinase and phosphatase), all the relevant quantities like 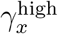, *A*, and *I* can be calculated.

Naively one might expect evolution to always drive systems to optimality due to natural selection, but genetic drift can play a significant competing role, allowing sub-optimal variants to flourish and even fix in a population [59]. To be specific, let us consider a unicellular organism that reproduces via binary fission, and two genetic variants of that organism that differ in the enzymatic parameters of a push-pull signaling loop: both variants achieve the same 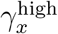 and *ℐ*, but one has *A > A*_min_ and one has *A* = *A*_min_. Let us denote the relative fitness of the sub-optimal versus the optimal type as 1 + *s*, defining a selection coefficient *s*. In other words the sub-optimal variant will have on average 1 + *s* offspring relative to the optimal one during the generation time of the optimal type. In the scenario described above, where the extra production does not confer any adaptive advantage and only imposes a metabolic cost, we will have *s <* 0, because the superfluous ATP consumption will lead to slower growth.

The magnitude of *s* determines the degree of selective pressure on the sub-optimal variant. The key quantity that sets the relevant scale for *s* is the effective population *N*_*e*_ of the organism, the size of an idealized population that exhibits the same changes in genetic diversity per generation due to drift as the actual population [58]. When *s <* 0 and |*s*| ≫ 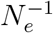, natural selection dominates drift, exponentially suppressing the probability of a sub-optimal mutant fixing in a population of optimal organisms. On the other hand if |*s*| ≪ 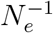, drift dominates, and the fixation probability of sub-optimal mutants is roughly the same as for a neutral (*s* = 0) mutation [60]. In this case it would be difficult to maintain optimality in a population over the long term. *N*_*e*_ for organisms is typically smaller than their actual population in the wild, and varies by several orders of magnitude among different classes: for unicellular species it can be as high as *∼* 10^9^ − 10^10^ in bacteria down to *∼* 10^6^ − 10^8^ in single-celled eukaryotes [32, 58]. (It becomes even smaller among higher eukaryotes, going down to *∼* 10^4^ in vertebrates.) The corresponding ranges for the “drift threshold” 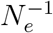 [32] are shown on the right in Fig. 5B.

The question then becomes: how do we estimate *s* and how does it compare to the relevant 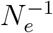 for the class of interest? For the case where a variant imposes metabolic costs but no adaptive advantage, there is a very useful relation that posits *s ∼* −*δC*_*T*_ */C*_*T*_ [32, 61, 62]. Here *C*_*T*_ is the total resting metabolic expenditure of an organism during a generation time, measured for example in units of P, where 1 P = one phosphate bond hydrolyzed (ATP or ATP equivalent consumed). *δC*_*T*_ is the extra expenditure incurred by the more costly mutant. This relation has already been used to explore selective pressures in yeast [62], unicellular prokaryotes and eukaryotes [32], and viral infections [63]. It was recently derived from first principles through a general bioenergetic growth model [33], where the relation was refined with a more accurate prefactor: *s ≈* − ln(*R*_*b*_)*δC*_*T*_ */C*_*T*_. Here *R*_*b*_ is the mean number of offspring per individual (i.e. *R*_*b*_ = 2 for binary fission).

The value of *C*_*T*_ can be readily estimated for single-celled organisms, where it scales roughly with cell volume [32, 33]. Given the 30 fL cell volume used in our calculations, and assuming a generation time (cell division time) *t*_*r*_ = 1 hr, we find *C*_*T*_ *≈* 7*×*10^11^ P (see details in the SI), comparable in magnitude to experimental estimates for yeast [32]. Since *δC*_*T*_ reflects the extra ATP consumed by the costly mutant (with consumption rate *A*) versus the optimal variant (rate *A*_min_) over one generation time, we can write *δC*_*T*_ = (*A* − *A*_min_)*t*_*r*_. We can thus calculate *s* for all the near-bandwidth *ℐ* = 1 bit parameter sets represented in Fig. 5A. The results for |*s*| versus 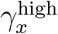 are plotted in Fig. 5B. Because increased ATP consumption is required to achieve larger bandwidths (as seen in Eq. (8)), the distribution of selective penalties |*s*| for being sub-optimal is pushed to larger values with greater 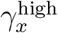. In other words, higher bandwidths make the energetic stakes more significant.

We can now rationalize why the yeast Pbs2/Hog1 loop might be close to optimality. The bandwidth for that system (indicated by a vertical dashed line in Fig. 5B) is near the higher end of the spectrum. Suboptimal parameter values that achieve approximately the same bandwidth at *ℐ* = 1 bit span a range of |*s*| values between 10^−8^ and 10^−4^. Given *N*_*e*_ = 10^6^ − 10^8^ for single-celled eukaryotes [32, 58], and estimates of *N*_*e*_ *≈* 10^7^ for wild yeast populations [64], these suboptimal systems likely have |*s*| near or above the drift threshold 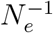. Thus we would expect yeast to be under evolutionary pressure to optimize the energy expenditures associated with the enzymatic loop.

## IV. DISCUSSION AND CONCLUSIONS

The kinase-phosphatase push-pull signaling network, which maintains a certain value of mutual information *ℐ* between input and output, incurs energetic costs in the form of ATP consumption. These costs have two related facets: (i) the free energy expenditure Δ*μ* for each hydrolysis reaction, and (ii) the number of such reactions *A* per unit time. Achieving empirical values like *ℐ* = 1 − 2 bits requires satisfying both aspects of the cost. There is a minimal price in terms of Δ*μ* to achieve any given *ℐ*, and this price increases if one demands either greater fidelity (larger *ℐ*) or the ability to process faster signals (larger *γ*_*x*_). Modern cells are more than willing to pay this part of the price, with Δ*μ* sufficiently high to meet the minimal requirements for any enzymatic parameter set that hits a target *ℐ* on the order of 1 bit. However, as the distributions in Fig. 3D-F illustrate, there are certainly options for signaling systems that work at similar fidelities under conditions of smaller Δ*μ*, the presumptive scenario earlier in evolutionary history. In all cases we require some degree of fine-tuning of enzymatic parameters: the higher the fidelity or frequency demands, the smaller the fraction of parameter space that satisfies them. This leaves vanishingly small room to achieve networks that operate at *ℐ* significantly larger than the known empirical range.

For particular parameter combinations the system is optimal, exhibiting the maximum possible bandwidth (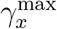 of Eq. (5)) with the minimal ATP consumption (*A*_min_ of Eq. (8)). These theoretical bounds can also be generalized to the case of time-delayed MI, giving the simple expression for the overall maximum MI shown in Eq. (6). Is such optimality widely realized in nature? Analyzing the selective pressures due to superfluous ATP expenditures indicates that this is a worthwhile question to pursue. We have already identified one near-optimal candidate in the yeast Hog1 signaling pathway. Based on the results of the previous section, we predict that the best place to look for others is among signaling pathways with high bandwidths, for example *∼* 10^−3^ − 10^−2^ s^−1^ at the extremes of the current biological distribution. Here the metabolic costs of being suboptimal would be significant for single-celled organisms.

More broadly, strong selective pressure on the costs of running signaling networks in single-celled organisms is likely to be a widespread phenomenon. To give another example, the expenditure of running the chemotaxis machinery in *E. coli* has been estimated to be about *∼* 10^7^ P per *∼* 1 hr cell cycle [2, 4]. Compared to a value of *C*_*T*_ *≈* 2 *×* 10^10^ P for *E. coli* [32, 33], we get an |*s*| *∼* 10^−4^, which is definitely significant for a bacterial population. We have barely begun to understand the kinds of optimization that such selective pressure has induced. Our approach readily generalizes beyond the kinase-phosphatase system, setting the stage for exploring these issues in a much wider array of biochemical networks.

## Data and code availability

The code for our analysis, along with the data used to generate the figures, is available at: https://github.com/hincz-lab/cell-signaling.

## Acknowledgments

We thank Shishir Adhikari for useful discussions. Parts of the numerical analysis were carried out using the High Performance Computing Resource in the Core Facility for Advanced Research Computing at Case Western Reserve University.

## Supplementary Information

### I. DERIVATION OF THE LOCAL DETAILED BALANCE RELATION

**FIG. S1.**
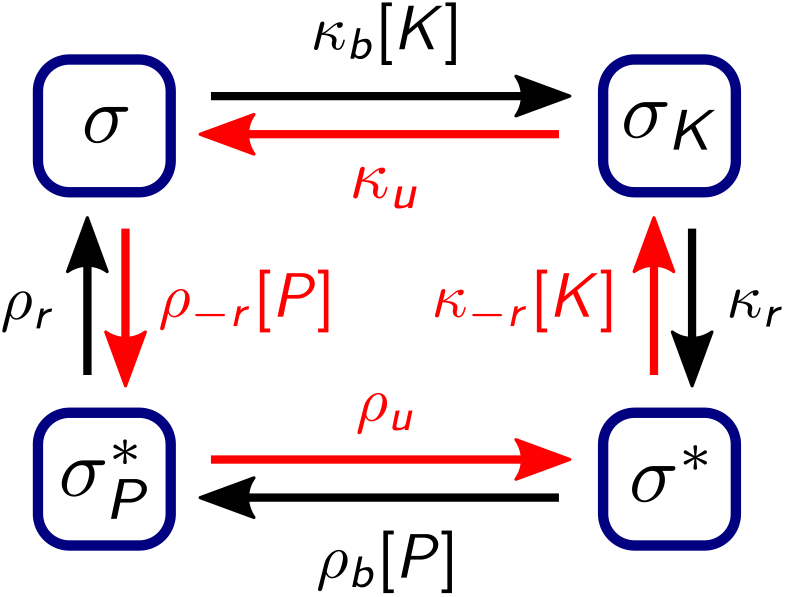
The enzymatic push-pull loop from the perspective of an individual substrate molecule. The protein can exist in one of four states: unmodified substrate (*σ*), bound to kinase (*σ*_*K*_), phos-phorylated (*σ*^*∗*^), and bound to phosphatase while phosphorylated 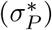. The forward (clockwise) transition rates between these states are indicated in black, while the reverse (counterclockwise) rates are in red.

To derive the local detailed balance relation of main text Eq. (1), it is convenient to focus on the reactions from the perspective of an individual substrate molecule [1]. A given molecule in our model can be in one of four states, indicated in Fig. S1 with corresponding forward and reverse transition rates. For example if the molecule is an unmodified substrate (state *σ*) it can transition to a kinase-bound substrate (state *σ*_*K*_) with rate *κ*_*b*_[*K*], proportional to the surrounding concentration [*K*] of kinase molecules. It can revert from *σ*_*K*_ to *σ* with rate *κ*_*u*_. The other transitions in Fig. S1 are defined analogously, with forward rates colored black and reverse rates in red. Local detailed balance entails that product of reverse rates divided by the product of forward rates is equal to exp(*β*Δ*G*), where Δ*G* is the free energy change of the system associated with a single forward traversal of the loop and *β* = (*k*_*B*_*T*)^−1^ [1]. Since after one loop from *σ* to *σ* the substrate is back in the same state (as well as the kinase and phosphatase), there is no contribution to Δ*G* from these molecules. However a single loop leads to the hydrolysis of a single molecule of ATP, so Δ*G* =−Δ*μ*, as defined in the main text. Putting everything together, the local detailed balance relation reads

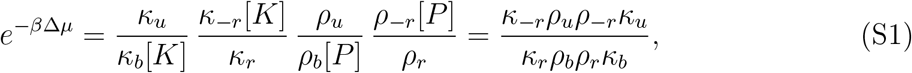

yielding main text Eq. (1).

### II. CHEMICAL LANGEVIN APPROACH FOR THE KINASE-PHOSPHATASE PUSH-PULL LOOP

In this section we derive the stationary state properties of the kinase-phosphatase push-pull loop via the chemical Langevin approximation. The derivation will follow analogously to Ref. [2], except here the system is more complicated due to the inclusion of reverse enzymatic reactions. The end goal will be a method to estimate the instantaneous mutual information *ℐ*, given by main text Eq. (4),

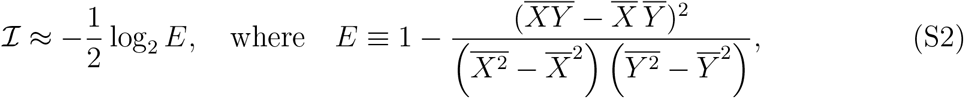

which requires evaluating the variances of the input and output, var 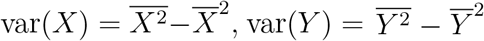, as well as the covariance cov 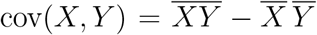. The quantity *E* here will be referred to as the “error” in signal propagation between input and output, and can be equivalently expressed as *E* = 1− *ρ*^2^, where *ρ* is the Pearson correlation coefficient between *X* and *Y*.

#### A. Dynamical equations

Our starting point is the full system of reactions for the enzymatic push-pull loop,

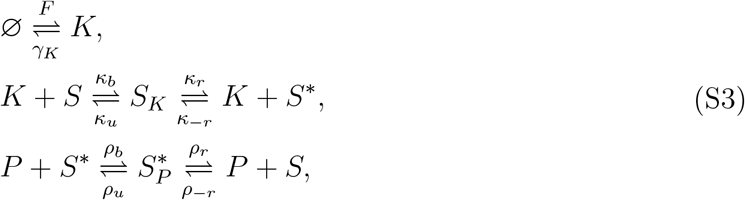

where ∅ represents the void (upstream deactivated kinase which does not enter into our model). The corresponding chemical Langevin equations [3] are given by:

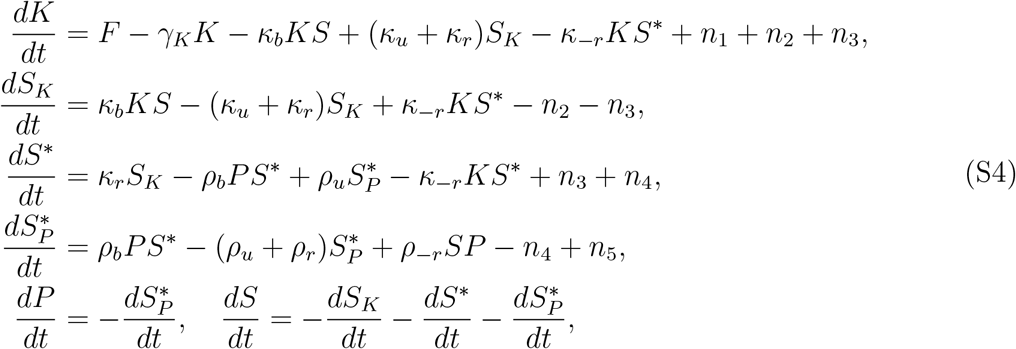

where the last line ensures that the total populations of free or bound phosphatase 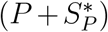 and free or bound substrate in all its forms 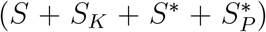 remain constant. The noise terms 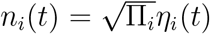 where *η*_*i*_(*t*) are Gaussian noise functions with zero mean and correlations 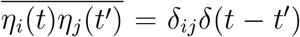. The five noise terms are associated with reactions in the system, and the corresponding prefactors represent the sum of the mean production (forward) and deactivation/unbinding (backward) contributions to each reaction:

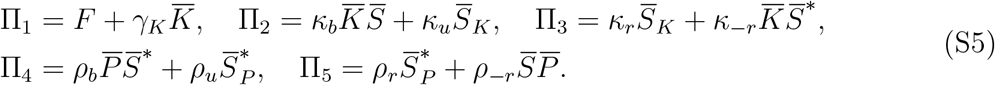

As described in the next section, we will be linearizing Eq. (S4), keeping terms up to first order in deviations from the stationary state values. In this linearized approach, the stationary state populations are given by:

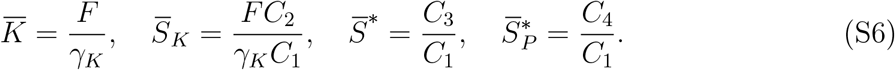

with the following definitions:

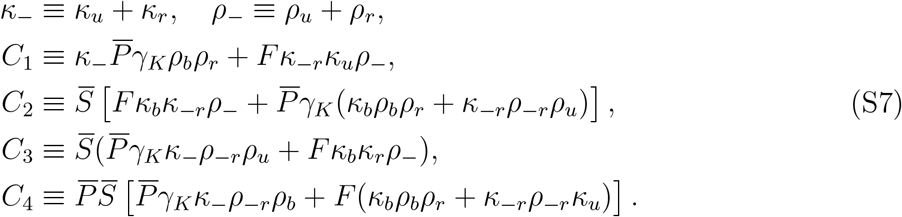

The input (total kinase) is *X* = *K* + *S*_*K*_ and the output (total activated substrate) is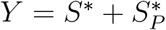, and hence Eq. (S6) can be used to calculate the stationary values 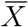 and 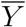.

#### B. Second moments

In order to calculate the variance and covariance of the input and output, we also need to know 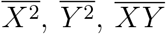. To estimate these quantities, the first step is to switch variables in Eq. (S4) to focus on deviations from the stationary state values: 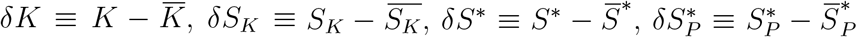. We can in turn rewrite these four variables in terms of the input and output deviations 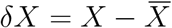 and 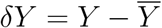 :

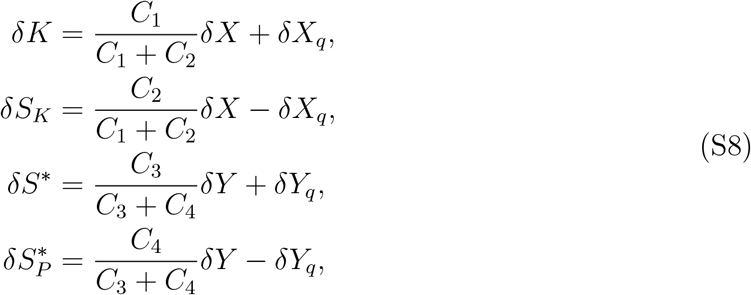

where we have introduced two additional auxiliary variables *δX*_*q*_ and *δY*_*q*_. Plugging Eq. (S8) into Eq. (S4), we simplify the system through linearization, ignoring any terms of second order or higher in the deviations. As demonstrated below in comparisons with kinetic Monte Carlo (KMC) simulations [4] of the original system, this linearized chemical Langevin approximation works well for our parameter ranges. Finally, we Fourier transform the linearized Eq. (S4), and the resulting system of equations takes the form

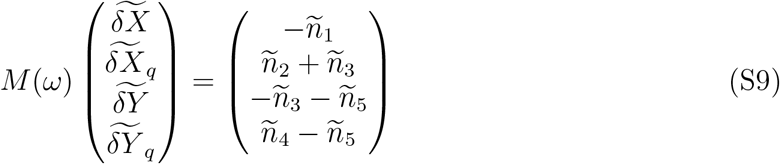

where 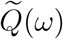 denotes the Fourier transform of quantity *Q*(*t*). The matrix *M* is given by:

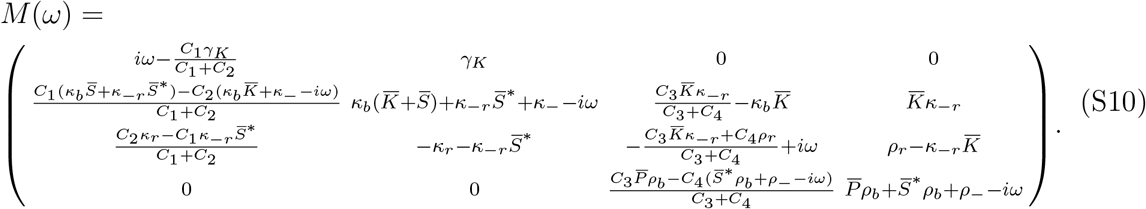

The Fourier-space system of equations Eq. (S9)-(S10) can be solved for 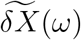 and 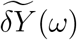. The expressions are complicated, but take the form of a linear combination of Fourier-space noise terms:

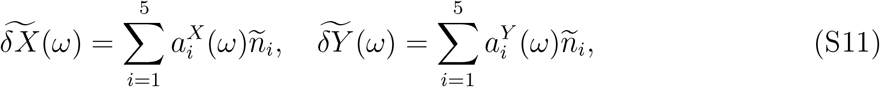

where 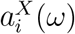 and 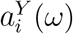 are some prefactors which can be expressed as rational functions of *ω*. The prefactors have the property 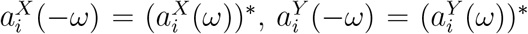. In Fourier space the correlations among the noise terms take the form 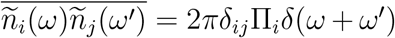. Hence we can calculate the input power spectral density (PSD) *P*_*X*_(*ω*), the output PSD *P*_*Y*_ (*ω*) and the cross PSD *P*_*XY*_ (*ω*), defined via

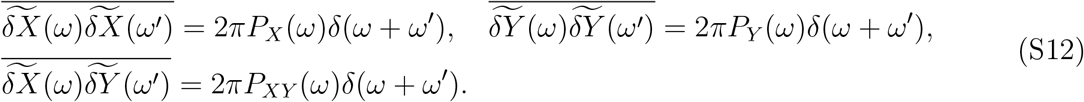

Plugging Eq. (S11) into Eq. (S12), we find expressions for the PSDs in terms of the prefactor functions:

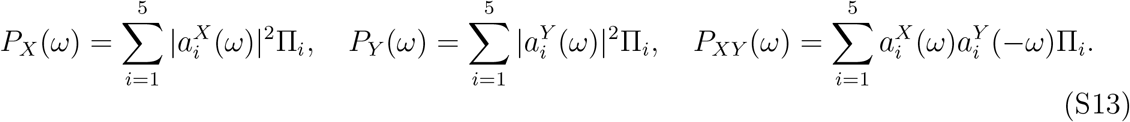

The final step is to calculate the second moments from integrals of the PSDs, using the inverse Fourier transform of Eq. (S13) evaluated at *t* = *t*^*i*^:

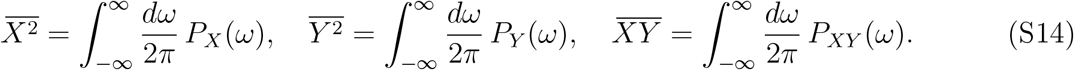

Given the explicit expressions for the prefactor functions in Eq. (S13) (which are available as part of the *Mathematica* notebooks in the Github repository associated with the manuscript), one can numerically evaluate the integrals in Eq. (S14) to get the moments.

#### C. Comparison to kinetic Monte Carlo simulations for mutual information

The chemical Langevin calculation of the second moments allows us to use Eq. (S2) to estimate the mutual information *ℐ*. We can then check whether this estimate is consistent with the results we would get from KMC simulations of the full system. Fig. S2 shows this comparison for two sample parameter sets drawn from the enzymatic parameter distribution described in Sec. IV. Since we are interested in exploring the full range of chemical potentials Δ*μ*, in each case we calculate *ℐ* varying the reverse-to-forward rate ratio *κ*_−*r*_*/κ*_*r*_, keeping all other parameters constant. Through main text Eq. (1), increasing *κ*_−*r*_*/κ*_*r*_ corresponds to decreasing the magnitude of Δ*μ*. At very large Δ*μ* (small *κ*_−*r*_*/κ*_*r*_) the *ℐ* curves saturate at the maximum possible mutual information for that parameter set, while at small Δ*μ* (large *κ*_−*r*_*/κ*_*r*_) the mutual information approaches zero, the equilibrium limit. Across the whole range we see that the chemical Langevin theoretical prediction is in close agreement with the KMC results.

### III. CHARACTERISTIC FREQUENCY *γ*_*x*_, GAIN *R*_0_, AND THE CONDITIONS FOR WIENER-KOLMOGOROV NOISE FILTER OPTIMALITY

#### A. Deriving the *γ*_*x*_ and *R*_0_ expressions in main text Eq. (2)

Since the effective frequency *γ*_*x*_ of the input and the gain *R*_0_ play central roles in the analysis, having simple closed form approximations for them [main text Eq. (2)] is useful. The original definitions of these two variables, as described in the main text, are as follows: (i) *γ*_*x*_ is related to the autocorrelation of input fluctuations, 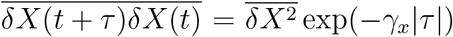 ; (ii) 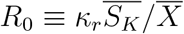 measures output production for a given level of input. As demonstrated in the next section, both of these can be calculated from KMC simulations (at significant computational expense for each different set of parameters). Alternatively, the chemical Langevin approximation of SI Sec. II can be used to derive somewhat cumbersome analytical expressions.

However the most convenient option is to take advantage of the meaning of *γ*_*x*_ and *R*_0_ in an effective, two-species description of the kinase-phosphatase reaction network. Imagine a system with an input species population *X*(*t*), output *Y* (*t*), and a simplified chemistry with only four reactions: production of input at rate *F*, deactivation of input at rate *γ*_*x*_*X*(*t*), production of output at rate *R*_0_*X*(*t*), and deactivation of output at rate *γ*_*y*_*Y* (*t*). In this two-species system the inverse input autocorrelation time is given by the deactivation rate parameter *γ*_*x*_, and the coefficient *R*_0_ in the output production rate is also the gain parameter. To relate this simplified model to the full reaction network of SI Sec. II, we compare analogous quantities in the simplified and full schemes. For example, let us take the mean input population 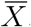. In the simplified scheme this is given by

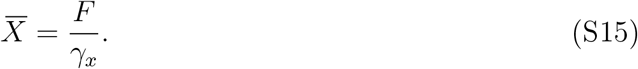

In the full network 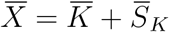 can be calculated from Eq. (S6) as

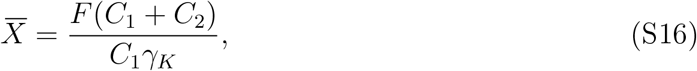

where the *C*_*i*_ are expressed in terms of full network parameters in Eq. (S7). Comparing Eqs. (S15) and (S16) we see that *γ*_*x*_ should be given by

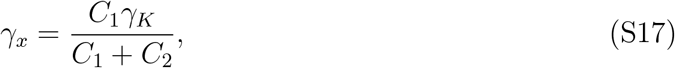

which is the first expression in main text Eq. (2). Similarly the mean production rate of the output in the simplified scheme is 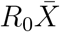. In the full system the mean output production is the average rate at which new phosphorylated substrate is produced via catalysis by the kinase-substrate complex,

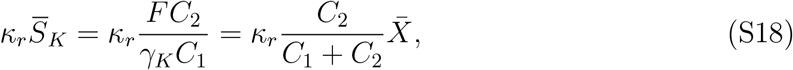

where we have again used Eqs. (S6)-(S7). Comparing Eq. (S18) to 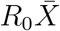, we see that *R*_0_should correspond to

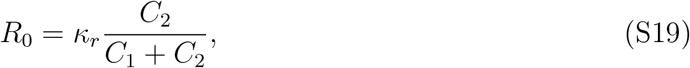

which is the second expression in main text Eq. (2).

**FIG. S2.**
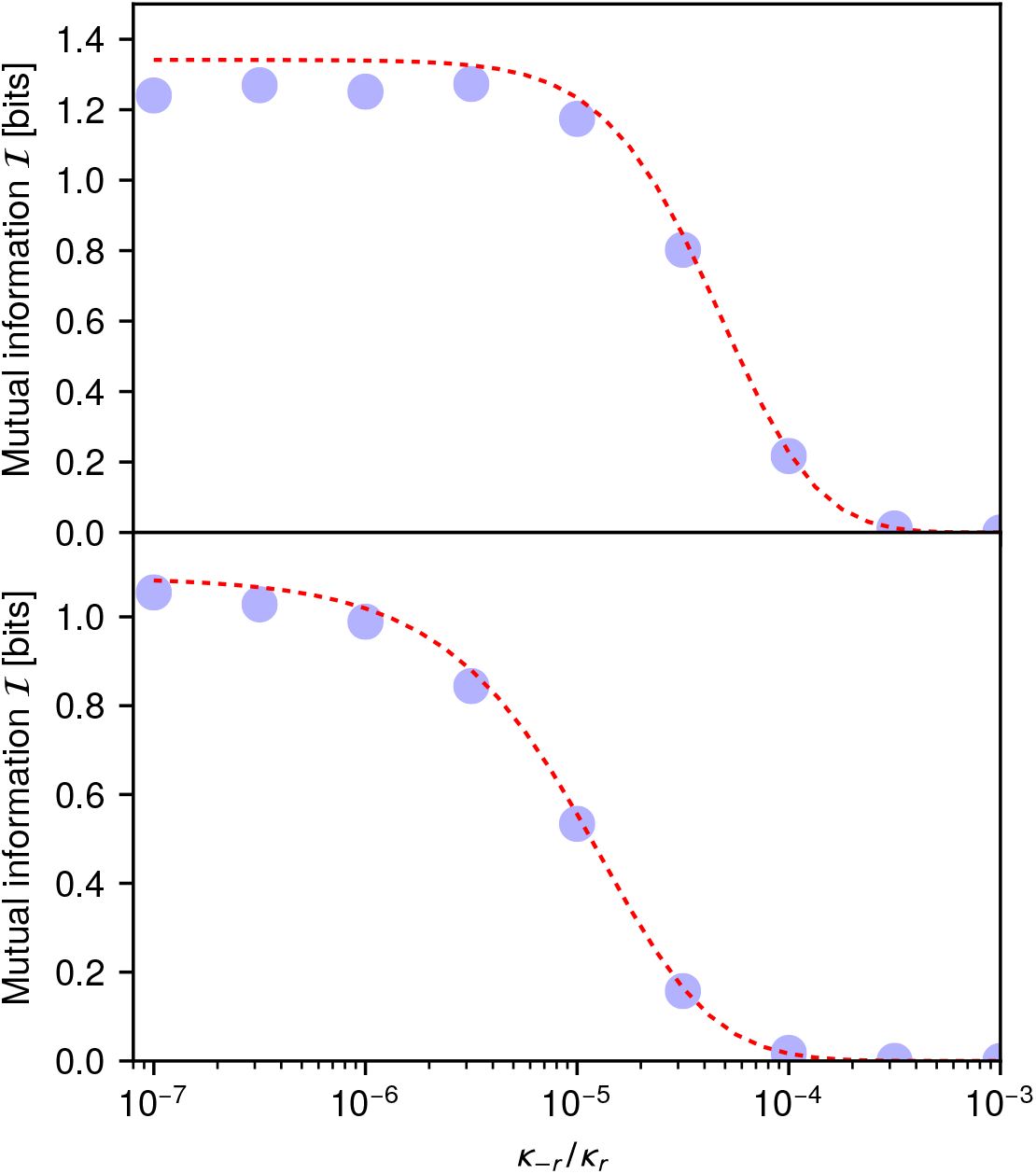
The mutual information ℐ for the enzymatic push-pull loop as a function of the reverse-forward rate ratio 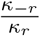. The predictions from the chemical Langevin approach (dashed line) are compared against the corresponding KMC simulation results (circles). The parameters sets are as follows (all units are s^−1^ except for the mean populations; molar units have been converted to populations by assuming a cell volume of 30 fL): (top) *κ*_*b*_ = 2.94 *×* 10^−6^, *ρ*_*b*_ = 3.68 *×* 10^−7^, *κ*_*u*_ = 1.58 *×* 10^−2^, *ρ*_*u*_ = 4.42 *×* 10^−4^, *κ*_*r*_ = 12.8, *ρ*_*r*_ = 1.34, *ρ*_−*r*_ = 2.50 *×* 10^−5^, *F* = 2.49 *×* 10^−3^, *γ*_*k*_ = 2.68 *×* 10^−5^, 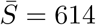 and 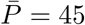 ; (bottom) *κ*_*b*_ = 2.32 *×* 10^−5^, *ρ*_*b*_ = 1.46 *×* 10^−4^, *κ*_*u*_ = 6.94 *×* 10^−2^, *ρ*_*u*_ = 5.48, *κ*_*r*_ = 0.994, *ρ*_*r*_ = 5.05 *×* 10^−2^, *ρ*_−*r*_ = 2.06 *×* 10^−8^, *F* = 2.46 *×* 10^−2^, *γ*_*k*_ = 2.65 *×* 10^−4^, 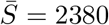 and 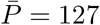.

#### B. Validation through kinetic Monte Carlo simulations

To verify that the expressions for *γ*_*x*_ and *R*_0_ derived above are good approximations, we ran KMC simulations for various parameter sets drawn at random from the enzymatic parameter distribution detailed in the SI Sec. IV. For each parameter set the simulation was run long enough after reaching the stationary state to collect sufficient statistics for both the mean population values and the input autocorrelation function. As described above, these allow us to calculate *γ*_*x*_ and *R*_0_. The simulation results are compared against the approximation from Eqs. (S17) and (S19) in Fig. S3. The agreement is excellent for both quantities, across the entire range of *γ*_*x*_ and *R*_0_ values. Thus we can confidently use the simple analytical expressions of Eqs. (S17) and (S19) to predict *γ*_*x*_ and *R*_0_ for any given parameter set.

#### C. Relating maximum bandwidth, minimum ATP consumption rate, and mutual information via Wiener-Kolmogorov optimal noise filter theory

One of the benefits of the approximate relation between the full system and the two-species model described in Sec. IIIA is that it allows us to use results from the two-species case to make predictions for the behavior of the kinase-phosphatase push-pull loop. The two-species model has been analyzed in detail in Refs. [2, 5], where it was shown to be able to map onto a Wiener-Kolmogorov optimal noise filter. The error *E* from Eq. (S2) for the two-species case can be evaluated in closed form as [2]:

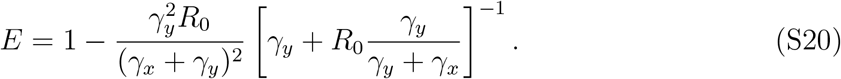

It achieves its minimum value (hence maximizing the mutual information *ℐ*) when the following condition is fulfilled:

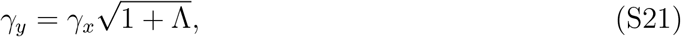

where Λ = *R*_0_*/γ*_*x*_. The corresponding minimum *E*, where the system behaves like an optimal Wiener-Kolmogorov (WK) noise filter is given by:

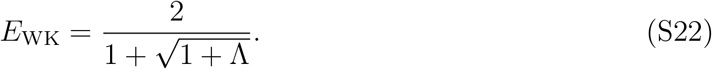

Interestingly, this remains the bound even if we generalize the output production term *R*_0_*X*(*t*) to be nonlinear in *X*(*t*) [2]. Using the relation between *E* and *ℐ* in Eq. (S2), we can translate the bound *E ≥ E*_WK_ into an equivalent statement that 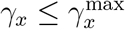 at a given value of mutual information *ℐ*. The value of 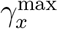 is shown in main text Eq. (5):

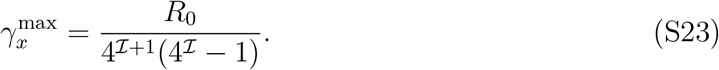

**FIG. S3.**
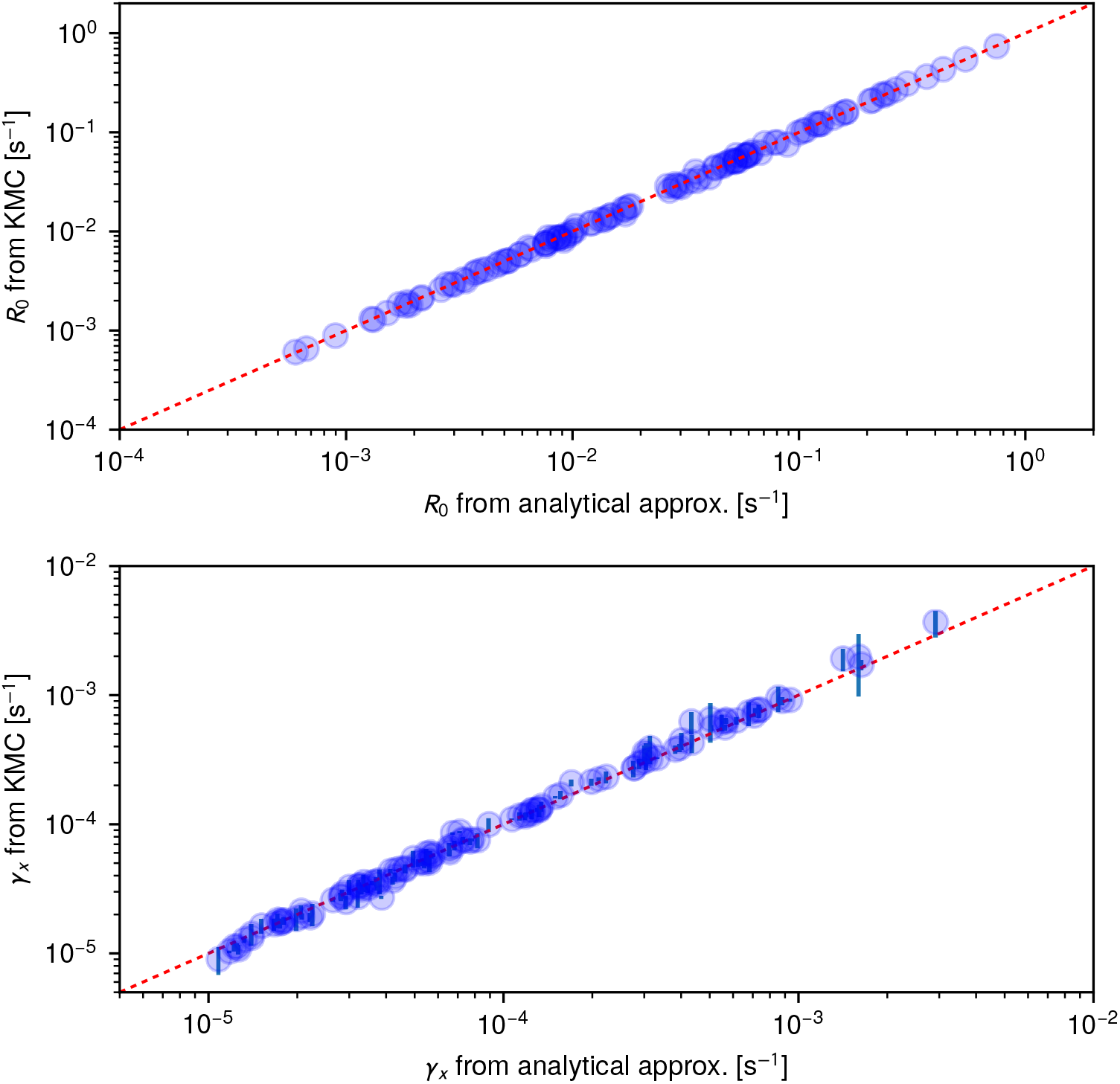
Comparison of the simple analytical approximations for *R*_0_ from Eq. (S19) (top) and *γ*_*x*_ from Eq. (S17) (bottom) versus KMC simulation results. Each point corresponds to a parameter set drawn randomly from the enzymatic parameter distribution described in SI Sec. IV. The red dashed line corresponds to perfect agreement. Error bars for *R*_0_ are smaller than the symbol size, and hence not indicated in the figure.

As shown in main text Figs. 3G-I, the above 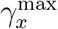 expression provides an excellent approximate upper bound on the 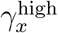 values calculated for the full enzymatic system. Even though the effective two-species model lacks reverse rates, it provides a useful tool for deriving this bound, since the maximum bandwidth is achieved when the reverse rates are negligible (large

As mentioned in the discussion around main text Eq. (7), the expression for 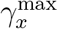 in Eq. (S23) also has an alternative interpretation. This gives the minimum production rate 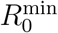 necessary to achieve mutual information *ℐ* at a certain bandwidth 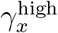 :

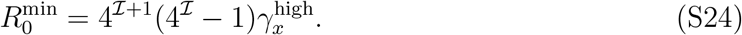

By relating *R*_0_ in turn to the ATP consumption rate 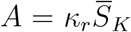, we can convert Eq. (S24) into an expression for the minimum necessary ATP consumption rate *A*_min_. To accomplish this, note that *A* can be rewritten as:

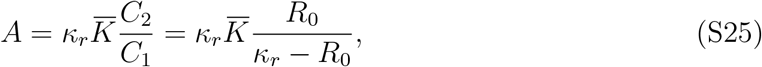

where we have used Eqs. (S6) and (S19). Finally, taking advantage of the fact that typically *κ*_*r*_ ≫ *R*_0_ for the parameter distributions of interest, we make the approximation 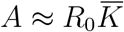. This allows us to derive main text Eq. (8):

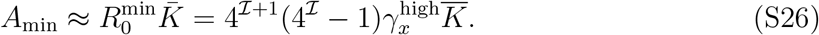

#### D. Optimality with respect to time-offset mutual information

The two-species optimality calculations of the previous section can be generalized to investigate mutual information *ℐ*_*α*_ between input *X*(*t*) and output *Y* (*t* + *α*) with a time offset *α*. Following the formalism of Ref. [5], the Wiener-Kolmogorov minimal error *E*_WK,*α*_ for arbitrary *α* is

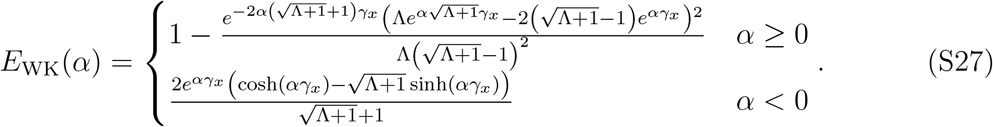

For *α* = 0 this expression reduces to *E*_WK_ from Eq. (S22). The associated mutual information ℐ_WK_(*α*) = − (1*/*2) ln_2_ *E*_WK_(*α*) varies with *α* in a manner qualitatively similar to main text Fig. 4A. It peaks at a value 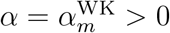 given by:

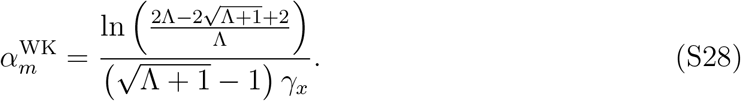

The mutual information 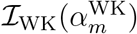 at the peak can be expressed as

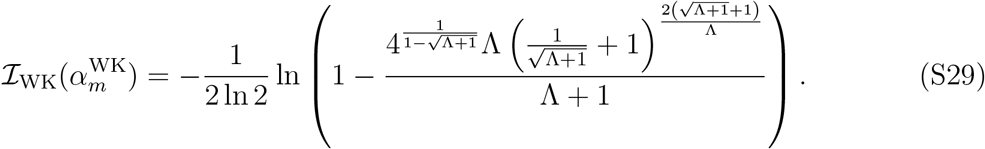

Since the expression in Eq. (S29) is somewhat unwieldy, we can simplify it using a close upper bound,

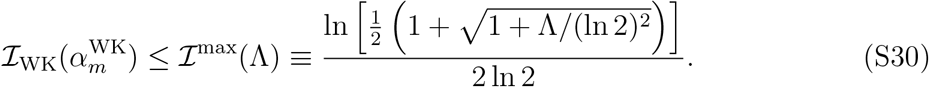

**TABLE S1.** Results of log-normal fits to various kinase/phosphatase enzymatic parameters. For each fit the mean log_10_ 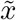 and standard deviation *σ*_*x*_ are listed. The top rows of the table correspond to individual fits to parameters collected from the PaxDb and Sabio-RK databases. The bottom rows show the results of a joint fit, described in the text of SI Sec. IV

| Parameter $x$ | Unit | $\log_{10} \tilde{x}$ | $\sigma_x$ | Data source |
| --- | --- | --- | --- | --- |
| *direct fits to database values:* |  |  |  |  |
| kinase/phosphatase concentrations [S], [P] | [M] | -7.93 | 0.84 | PaxDb [8] |
| Michaelis constants $K_M^{\text{kin}}, K_M^{\text{pho}}$ | [M] | -4.26 | 1.21 | Sabio-RK [9] |
| specificity ratios $\kappa_r/K_M^{\text{kin}}, \rho_r/K_M^{\text{pho}}$ | [M<sup>-1</sup> s<sup>-1</sup>] | 3.86 | 1.19 | Sabio-RK [9] |
| reaction rates $\kappa_r, \rho_r$ | [s<sup>-1</sup>] | -0.04 | 1.16 | Sabio-RK [9] |
| *results of joint fitting:* |  |  |  |  |
| reaction rates $\kappa_r, \rho_r$ | [s<sup>-1</sup>] | -0.06 | 1.18 | joint fit |
| binding rates $\kappa_b, \rho_b$ | [M<sup>-1</sup> s<sup>-1</sup>] | 3.94 | 1.12 | joint fit |
| dissociation constants $K_D^{\text{kin}}, K_D^{\text{pho}}$ | [M] | -7.00 | 1.31 | joint fit |

The bound ℐ^max^(Λ) is never more than ≈ 0.04 bits above 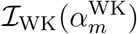 for any Λ, and converges to 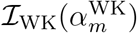 in the limit Λ → ∞. Hence for all practical purposes we can use Eq. (S30) in place of Eq. (S29) to estimate the bound on the mutual information, as we do in main text Eq. (6).

### IV. ENZYMATIC PARAMETER DISTRIBUTION

Earlier surveys of enzymatic kinetic parameters in Refs. [6, 7], over broader classes than just kinases and phosphatases, showed that their distributions could be approximately described by log-normal distributions. For a given parameter *x*, we will denote this as 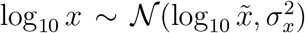 or in other words that the base-10 logarithm of *x* is distributed according to a normal distribution with mean 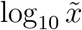 and standard deviation *σ*_*x*_. The value 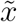 is the median of the resulting log-normal distribution for *x*.

For our work the focus is on kinases and phosphatases, and we are interested in looking at the push-pull loop signaling behavior over the entire distribution of biologically plausible parameters. The parameter data we collected, summarized in the histograms of main text Fig. 2, had far more representation of kinases than phosphatases, which is a well known limitation of the existing experimental literature. Despite this sampling issue, the orders of magnitude spanned by phosphatase parameters were comparable to those of the kinases. For each parameter type, we thus decided to fit both types of enzyme with a single overall distribution, based on pooling of all the available kinase and phosphatase data together. The data available from the databases took the forms listed below (all raw data and the files used to process it are included in the Github repository associated with the manuscript). The mean 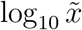 and standard deviation *σ*_*x*_ values from the log-normal fits for the different parameter classes are listed in the first four rows of Table S1.

Enzymatic data:

- Mean substrate [*S*] and phosphatase [*P*] concentrations, where the substrate is taken to be a kinase [main text Fig. 2A]. These numbers were derived from the PaxDb protein abundance database [8], taking advantage of UnitProt gene ontology associations to focus on just kinases and phosphatases in signal transduction pathways [10]. Each PaxDb data entry is in terms of ppm (parts per million) of abundance, relative to the total number of proteins in the cell. To convert from ppm to molar concentrations, we looked at data from human cells (which had the best representation in the database), and used the estimated total concentration of 2.7*×*10^6^ proteins per *μ*m^3^ for human cells [11]. The latter concentration corresponds to 4.48*×*10^−3^ M. If *y* is the abundance in ppm units, then 4.48(*y/*10^6^) *×*10^−3^ M is the corresponding molar concentration. Note that total concentrations are very similar across many different types of species [11], so there should not be a strong species-dependence in the analysis. For example the same analysis in mouse cells rather than human ones yields quantitatively similar results: a mean kinase/phosphatase concentration 10^−8.31^ M (versus 10^−7.93^ M in human cells), and a log-normal standard deviation of 1.03 (versus 0.84 in human cells).
- Reaction parameters [main text Fig. 2B-D]. These values were taken from the Sabio-RK database [9], where they were most often available in the following forms: Michaelis constants 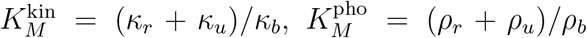 for the kinase/phosphatase (main text Fig. 2B), the corresponding specificity ratios 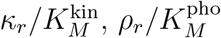 (main text Fig. 2C), and the reaction rates *κ*_*r*_ and *ρ*_*r*_ (main text Fig. 2D). The resulting distributions were entirely consistent (though slightly narrower) with the distributions for the same parameter types analyzed in Ref. [6], which considered all enzymes (not just kinases and phosphatases).

Note that the six reaction parameter types that were collected from the Sabio-RK database 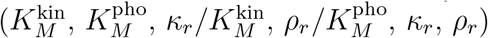 are not directly in the form that we need to calculate push-pull loop signaling properties. For the latter we would like to know (*κ*_*b*_, *ρ*_*b*_, *κ*_*u*_, *ρ*_*u*_, *κ*_*r*_, *ρ*_*r*_), or equivalently 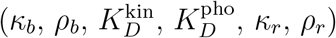. Here the dissociation constants are defined as 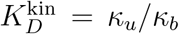 and 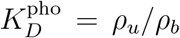. Let us denote the parameter vector 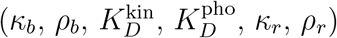 as **v**, with components *v*_*α*_, *α* = 1, …, 6. We would like to find a joint distribution for **v** that is self-consistent with the individual log-normal distributions for the alternative parameter types fitted directly from the database values (first 4 rows of Table S1). We will assume the simplest form for the joint distribution Φ: a product of individual log-normal distributions for each parameter *v*_*α*_, with median values 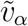 and standard deviations *σ*_*α*_:

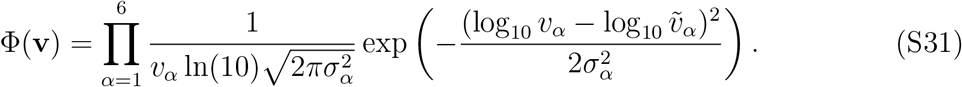

Note that the *v*_*α*_ ln(10) term in the denominator of the prefactor comes from the Jacobian due to the variable change between log_10_ *v*_*α*_ and *v*_*α*_. This ensures that the probability is properly normalized: 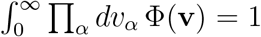. As explained above, kinase and phosphatase parameters are assumed to be drawn from the same distributions, so we enforce that 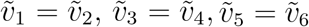, and analogously for the standard deviations *σ*_*α*_. This leaves six distinct values that determine the distribution: 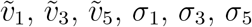.

To estimate these six distribution parameters, we use the following iterative numerical fitting procedure. We start with a guess for 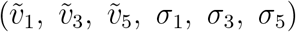 and then draw 10^4^ parameter sets **v** from the resulting distribution Φ(**v**). For each parameter set we can calculate the alternative parameter types 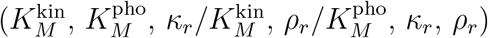. We then fit the resulting 10^4^ values for these alternative types to individual log-normal distributions, and compare the means and standard deviations to the empirical results in the top half of Table S1. The sum of the relative absolute errors between the new joint fit values and the empirical results for the means / standard deviations is our overall goodness-of-fit measure. We perturb our guess for 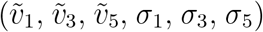 and accept the perturbation if it improves the goodness-of-fit. This procedure is iterated until convergence. The results of this joint fit are shown in the bottom half of Table S1. The joint fit predictions for the binding rate (*κ*_*b*_, *ρ*_*b*_) and dissociation constant 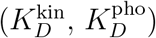 distributions are consistent with earlier estimates of these parameters in specific kinase/phosphatase systems [12]. As another consistency check, the joint fit distribution for the reaction rates (*κ*_*r*_, *ρ*_*r*_) is nearly identical to the individual empirical fit based on the Sabio-RK database values.

Finally we note that the simple joint distribution Φ(**v**) in Eq. (S31) is by construction too broad: it may produce the correct marginal distributions for quantities collected from the Sabio-RK database, but it ignores any correlations between those individual parameters that may be present in natural systems. Estimating these correlations from the existing database entries is quite challenging, because relatively few entries have a complete list of all the parameters of interest. Hence, as explained in the main text, we take Φ(**v**) to be effectively a superset: it should contain the true, presumably narrower, biological distribution plus parameter sets that are less likely to be observed in nature. A convenient aspect of this interpretation is that any collective conclusion we draw from the entire distribution Φ(**v**) should also be true for the subset of biological parameters. Moreover we can thus explore a larger design space (potentially available for evolution) than what we currently observe in modern biological systems.

### V. RESULTS FOR ALTERNATIVE INPUT KINASE CONCENTRATIONS

The results in main text Fig. 3D-F were for a mean input kinase concentration [*K*] = 5 nM. In Fig. S4 we show the analogous results for two different choices: [*K*] = 0.5 nM (left column) and [*K*] = 50 nM (right column). The main conclusions remain unchanged: the physiological Δ*μ* range (highlighted in pink) is always just above the upper edge of the 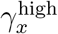 cloud, and the number of available parameter sets decreases rapidly as the mutual information *ℐ* is increased.

### VI. ANALYSIS OF THE PBS2-HOG1 PUSH-PULL LOOP IN YEAST

To illustrate our theoretical framework in a concrete biological example, let us consider a kinase-phosphatase loop from one of the most extensively studied signaling pathways: the Hog1 mitogen-activated protein kinase (MAPK) pathway that allows yeast to adapt to extracellular osmotic changes [13–15]. We will focus in particular on the final portion of the pathway, where the active (phosphorylated) kinase Pbs2pp catalyzes the conversion of inactive Hog1 into phosphorylated Hog1pp. The latter protein is interchanged quickly between cytoplasm and nucleus, where it regulates a variety of responses to osmotic stress. Hog1pp is dephosphorylated by a combination of phosphatases Ptp2 (mainly in the nucleus) and Ptp3 in the cytoplasm [16]. Thus Pbs2pp will play the role of *K* in our model, Hog1 will be *S*, Hog1pp will be *S*^*∗*^, and Ptp2/Ptp3 will be *P*. To parameterize our model, we start with a more detailed theoretical description of the entire pathway developed by Zi *et al*. [13]. A key appeal of this work is that its parameters were carefully fit to extensive experimental data from yeast cells exposed to different time series of external salt shocks in microfluidic experiments [14]. However since the parameters of Zi *et al*. are not expressed in the same form as the enzymatic reaction rates of our model, we do have to convert from their framework to ours, as described below.

**FIG. S4.**
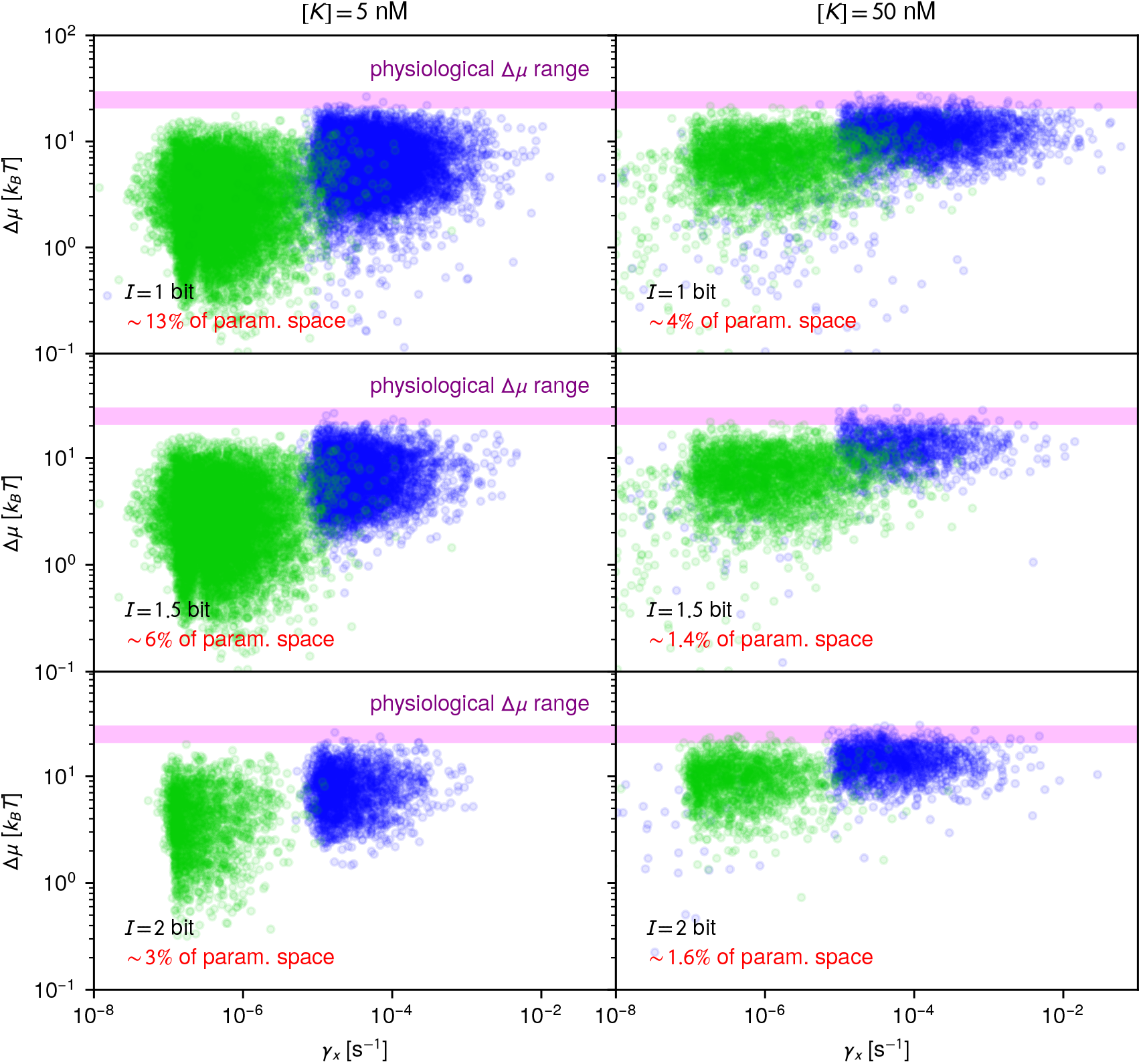
Analogous to main Figure 3D-F, except for input kinase concentration [*K*] = 5 nM (left column) and 50 nM (right column). The rows correspond to mutual information *ℐ* = 1, 1.5, and 2 bits respectively. The probabilities of successfully drawing a parameter set that achieves the specified *ℐ* value are shown in red in panel.

#### A. Parameter estimation based on earlier literature

Ref. [13] explicitly distinguishes between the concentration of Hog1 and Hog1pp in the cytoplasm and nucleus, denoted with c and n superscripts respectively: [Hog1^c^], [Hog1^n^], [Hog1pp^c^], [Hog1pp^n^]. If we are interested in the average concentrations overall, we can denote these as:

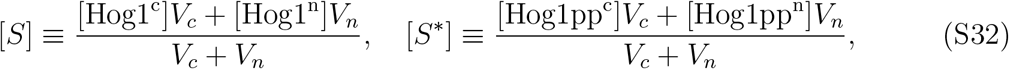

where *V*_*c*_ and *V*_*n*_ are the volumes of the cytoplasm and nucleus respectively, taken to have a ratio of *V*_*n*_*/V*_*c*_ = 0.14 [13]. Eq. (S32) also implies:

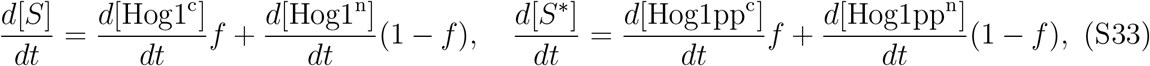

where *f* = *V*_*c*_*/*(*V*_*c*_ + *V*_*n*_) = 0.88. As a simplification of Eq. (S32), we note in Ref. [13] import and export of the Hog1 proteins is fast relative to other reactions, and for a given input level the system rapidly reaches a stationary state with [Hog1^n^]*≈*[Hog1^c^]*≈* [*S*], [Hog1pp^n^]*≈*[Hog1pp^c^]*≈* [*S*^*∗*^].

We can now look at individual reactions that contribute to the time derivatives on the right-hand sides of Eq. (S33) and find their analogues in our model. For example the phosphorylation step that converts Hog1^c^ to Hog1pp^cs^ is expressed in Ref. [13] as an effective second order reaction of the form 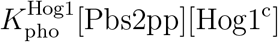, with rate constant 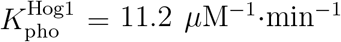. This contributes positively to *d*[Hog1pp^c^]*/dt* and with a minus sign to *d*[Hog1^c^]*/dt*, and so leads to contributions magnitude 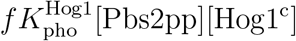 to the righthand sides of Eq. (S33). Note that even though activation of Hog1 is actually a double phosphorylation (of a threonine and tyrosine residue), the entire process in this case can be well approximated through a single rate constant.

In our model the conversion of *S* to *S*^*∗*^ occurs through the intermediate state *S*_*K*_. However if we want to compare to the phosphorylation step of Ref. [13] in order to match parameters, we can look at the deterministic contribution to the dynamics (ignoring fluctuations) in the Michaelis-Menten approximation for enzyme kinetics [17]. In this picture the phosphorylation reaction contributes to *d*[*S*]*/dt* and *d*[*S*^*∗*^]*/dt* through a term of magnitude 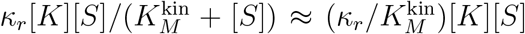, where the last simplification is valid when 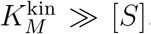. If we compare 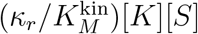 to 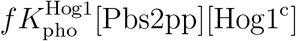, noting that [*K*] = [Pbs2pp] and [*S*] *≈* [Hog1^c^], we can make the following identification:

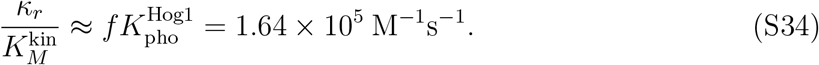

The dephosphorylation steps in Ref. [13] are modeled as two pseudo-first-order reactions: conversion of Hog1pp^c^ to Hog1^c^ with rate 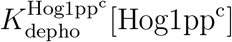, and the conversion of Hog1pp^n^ to Hog1^n^ with rate 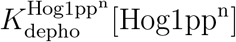. The pseudo-first-order rate constants are given by: 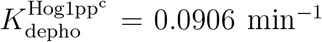 and 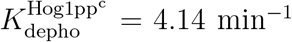. These reactions will lead to contributions of magnitude 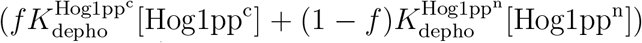 to the right-hand sides of Eq. (S33). In our model (using a similar Michaelis-Menten approximation to the one described above, with 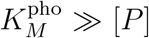), the analogous expression for dephosphorylation is effectively a second-order reaction with rate 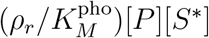. Comparison of the two expressions, using the approximation [Hog1pp^n^]*≈*[Hog1pp^c^]*≈* [*S*^*∗*^], leads to the identification:

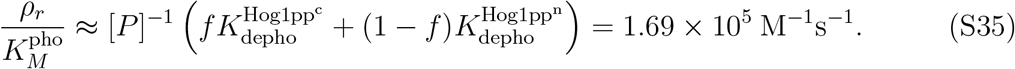

**TABLE S2.** Summary of parameters for the yeast Pbs2/Hog1 system estimated from earlier literature.

| Parameter | Value | Data source |
| --- | --- | --- |
| substrate concentration $[S]$ | $0.38 \mu\text{M}$ | Hog1 abundance from Ref. [18] |
| phosphatase concentration $[P]$ | $0.058 \mu\text{M}$ | Ptp2/Ptp3 abundance from Ref. [18] |
| kinase specificity $\rho_r/K_M^{\text{kin}}$ | $1.64 \times 10^5 \text{ M}^{-1}\text{s}^{-1}$ | analysis of Ref. [13] model fit to experiments of Ref. [14] |
| phosphatase specificity $\rho_r/K_M^{\text{pho}}$ | $1.69 \times 10^5 \text{ M}^{-1}\text{s}^{-1}$ | analysis of Ref. [13] model fit to experiments of Ref. [14] |

Here we set [*P*] = 0.058 *μ*M as an average measure of phosphatase concentrations, to facilitate the conversion from pseudo-first-order to second-order rate constants. The value of [*P*] is based on estimates of the concentrations of the two phosphatases in yeast from Ref. [18]: 0.049 *μ*M for Ptp3 in the cytoplasm, and 0.067 *μ*M for Ptp2 in the nucleus, where we have used *V*_*c*_ = *f* (*V*_*c*_ + *V*_*n*_), *V*_*n*_ = (1 − *f*)(*V*_*c*_ + *V*_*n*_) and *V*_*c*_ + *V*_*n*_ ≈ 30 fL [13, 19] to convert from populations to concentrations. Since the concentrations were of similar scale, we let [*P*] be the mean of the two values.

As a consistency check to make sure the final estimates of the specificity ratios 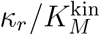 and 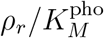 in Eqs. (S34)-(S35) are biologically plausible, we can compare them with the distribution of these ratios among kinases/phosphatases from the Sabio-RK database in main text Fig. 2C. The values for the Hog1/Pbs2 system are not unusual, and lie near the higher end of the range, at about the 0.87 quantile. The final parameter value we can estimate from the literature is the mean Hog1 concentration [*S*] = 0.38 *μ*M, based on the abundance reported in Ref. [18].

#### B. Estimation of remaining parameters

Based on the above analysis, we have estimates for four quantities in the Pbs2/Hog1 system drawn from the earlier literature: 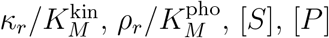. These are summarized in Table S2. The relationship of the enzymatic reaction/binding/unbinding rate parameters to the estimated values then takes the form:

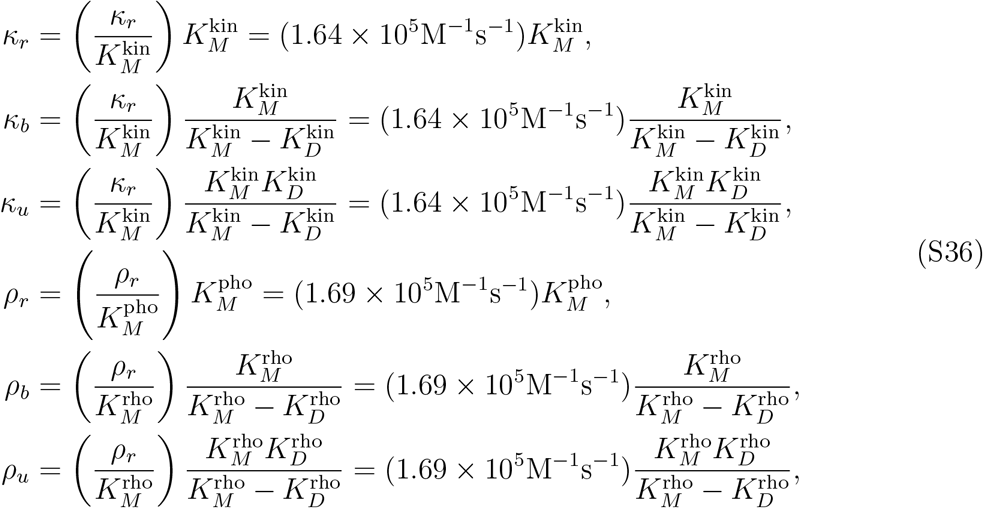

The above parameters depend on the values of 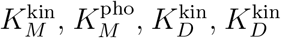. While we do not know what these are for the Pbs2/Hog1 system, we can draw their values from the corresponding empirical log-normal distributions described in Table S1. By repeating the draw many times, we can check how our final optimality analysis (see below) depends on the precise values of the unknown parameters. As it turns out the dependence of *R*_0_, 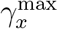 and 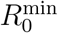 on the unknown values is quite weak, and we will be able to make robust estimates for these quantities. In the cases of 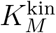 and 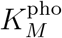, we constrain the random draw from their log-normal distributions to enforce 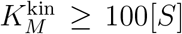 and 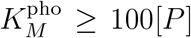. This ensures self-consistency with the assumptions 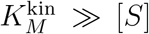 and 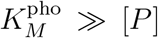, which were used in the previous subsection to match the form of the phosphorylation / dephosphorylation reactions between Ref. [13] and our model. The final two parameters are the reverse reaction rates *κ*_−*r*_ and *ρ*_−*r*_. Since we do not have any experimental estimates of these for the Pbs2/Hog1 system, we assume that the physiological value of Δ*μ* in yeast (around 21 *k*_*B*_*T* [19]) is sufficiently high that *κ*_−*r*_ and *ρ*_−*r*_ are negligible under normal conditions.

#### C. Bandwidth and gain

Given the parameter estimation procedure described above, we can calculate 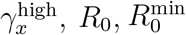 for each draw of the unknown parameters. The results remain within a narrow distribution, relatively insensitive to the values of the unknown parameters. The mean and standard deviations for 50 draws are: 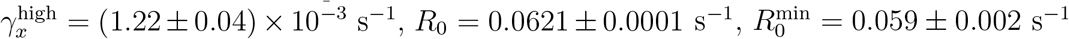

### VII. ESTIMATION OF TOTAL RESTING METABOLIC EXPENDITURE

For single-celled organisms, the total resting metabolic expenditure *C*_*T*_ can be estimated by the approach outlined in Ref. [20]. *C*_*T*_ has two contributions: *C*_*T*_ = *C*_*G*_ + *t*_*r*_*C*_*M*_. Here *C*_*G*_ is the expenditure involved in growth during one generation time *t*_*r*_, and *C*_*M*_ is the maintenance cost per unit time. Using a large collection of metabolic data from Ref. [21], covering both prokaryotes and single-celled eukaryotes, one can observe that both *C*_*M*_ and *C*_*T*_ scale approximately linearly with cell volume *V*, agreeing with the prediction of the bioenergetic growth model of Ref. [20]. The expression for *C*_*T*_ based on the results of these linear fits is [20]:

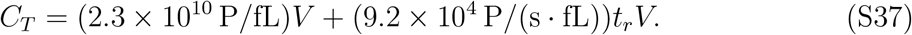

where the unit P corresponds to the hydrolysis of a phosphate bond (i.e. the consumption of one ATP or ATP equivalent). Using the main text values of *V* = 30 fL and *t*_*r*_ = 3600 s, we get *C*_*T*_ = 7.0 *×* 10^11^ P.

